# Genome-scale mapping of variant, enhancer and gene function in primary human CD4+ T cells

**DOI:** 10.64898/2026.03.09.710372

**Authors:** Dewi P. I. Moonen, Annique Claringbould, Andreas R. Gschwind, Stefan Schrod, Jana Braunger, Claudia Feng, Benedikt Rauscher, Jia Yi, Shirley Z. Bi, Yves Matthess, Manuel Kaulich, Ricelle A. Acob, Aruna Ayer, Jesse M. Engreitz, Britta Velten, Oliver Stegle, Gosia Trynka, Judith B. Zaugg, Daniel Schraivogel, Lars M. Steinmetz

## Abstract

CD4+ T cells harbor a disproportionate enrichment of immune disease risk loci and represent the primary cellular context for immune disease biology, yet the genes and regulatory programs these variants affect remain largely unknown. We combined targeted Perturb-seq of 1,032 *cis*-regulatory elements (CREs) overlapping 4,724 variants across 14 immune diseases with genome-wide Perturb-seq of all expressed genes in primary human CD4+ T cells, spanning 4.1 million cells. We identified 626 CRE-gene pairs, and connected CRE targets to downstream regulatory cascades. At the *TYK2* and *DEXI*/CLEC16A loci, we resolved target genes and linked noncoding variants to inflammatory and metabolic programs. Across diseases, we revealed that dispersed variants converged on shared and disease-specific programs. Our work provides a framework for tracing variant-to-CRE-to-gene-to-network in disease-relevant primary cells.

## Introduction

Genome-wide association studies (GWAS) have identified thousands of genetic variants associated with complex diseases, yet translating these discoveries into mechanistic understanding remains challenging. Over 60% of disease-associated variants reside in *cis*-regulatory elements (CREs), such as enhancers and silencers^1,2^. However, CRE-gene relationships cannot be reliably inferred from sequence or genomic context, and CRE activity is highly cell type- and state-dependent^3–5^. The central role of CREs in mediating disease risk is particularly evident in the immune system, where CD4+ T cells harbor a disproportionate enrichment of immune disease risk loci within chromatin regions that become specifically accessible during early T cell activation^6–8^. Despite this knowledge, only a handful of loci have been mechanistically dissected to identify the causal variant, target genes, and relevant cellular context^9^.

Several approaches have emerged for connecting noncoding variants and CREs to their target genes. Experimental approaches include massively parallel reporter assays^10,11^, chromatin conformation capture^12^, expression quantitative trait locus (eQTL) mapping^13^, and perturbation-based methods with single-cell transcriptomics^14–17^. While eQTL studies associate natural genetic variation to gene expression in many tissue contexts^18,19^, perturbation approaches (e.g. CRISPR(i/a) screens) provide direct experimental evidence but genome-wide studies are still uncommon and limited to few cellular contexts. In addition, these methods often capture distinct sets of genes to be regulated^20^. Computational approaches such as ENCODE-rE2G^21^ and enhancer-based gene regulatory networks (GRNs) allow genome-wide predictions, but GRNs are not transferable between cell types and machine learning models have not been validated outside the cell types in which training data has been generated.

Here, we present a systematic functional characterization of immune disease-associated CREs at genome scale in activated primary human CD4+ T cells, solving the above-mentioned limitations in scalability and cell type specificity. We performed sensitive targeted perturbation sequencing (TAP-seq) and genome-wide Perturb-seq in over 4.1 million cells, enabling network inference in the relevant biological context and providing a foundation for mechanistic interpretation of immune disease risk loci.

## Results

### Study design to map variant-to-CRE, CRE-to-gene, and gene-to-network

We implemented a multi-stage strategy combining *in silico* predictions and large-scale experimental testing to systematically link immune-disease variants to CREs, genes, and regulatory networks in primary human CD4+ T cells (Fig. 1A). First, we overlapped GWAS variants from 14 immune diseases with CD4+ T cell specific regulatory regions to prioritize 1,032 disease relevant CREs (variant-to-CRE). Second, we perturbed these CREs using CRISPRi and measured effects on all candidate target genes using targeted Perturb-seq (TAP-seq) ^15^ in over 1.5 million cells (CRE-to-gene, “CRE screen”). Third, we mapped downstream regulatory networks by genome-wide Perturb-seq of all expressed coding genes, yielding transcriptome-wide responses across 2.6 million cells (gene-to-network, “promoter screen”). In both screens, cells from the same three healthy donors (table S1) were profiled three days post re-stimulation as immune disease risk loci are specifically active during T cell activation. Each stage is detailed below.

**Fig. 1:**
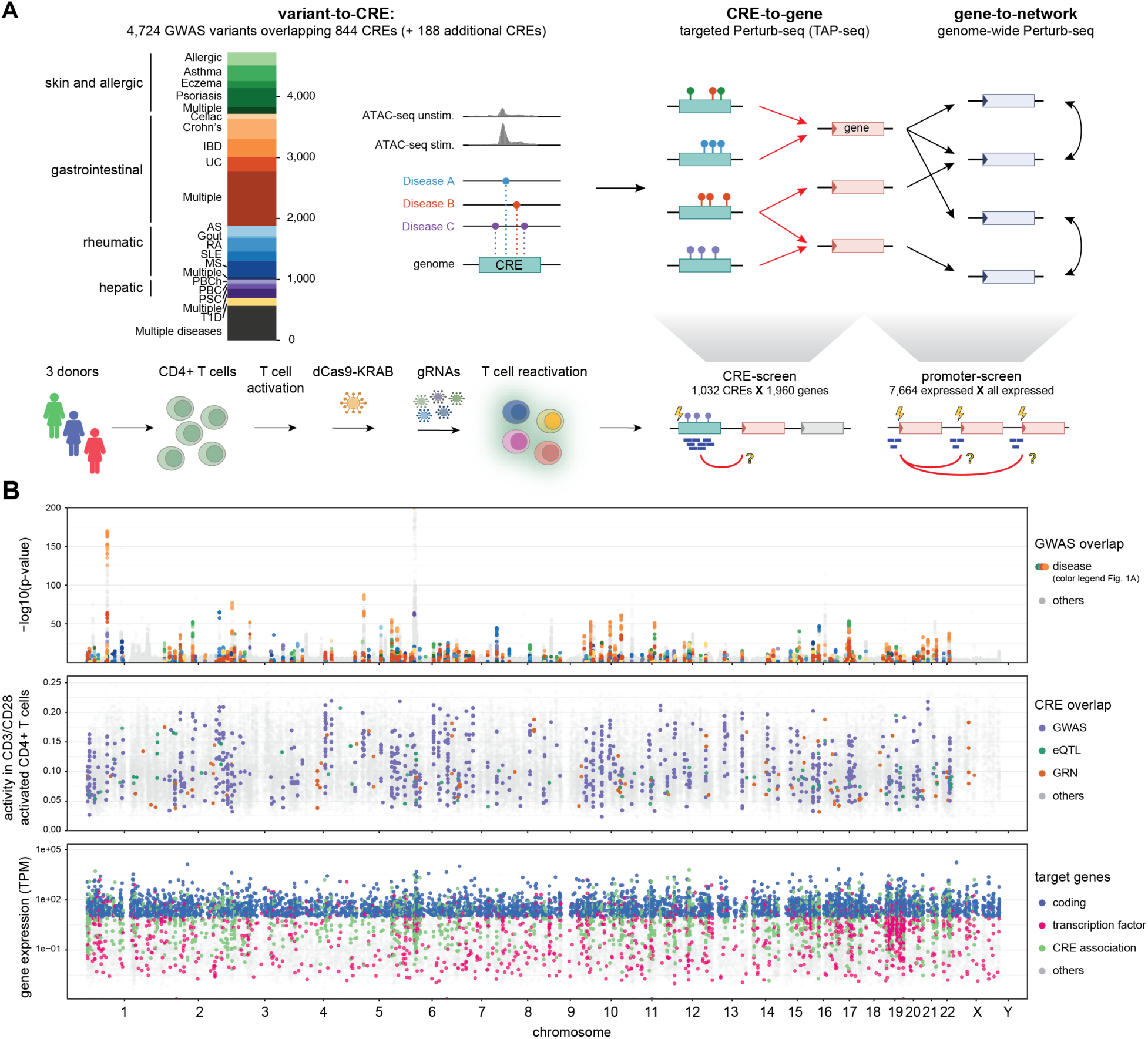
Study overview. (**A**) Study design from immune disease variants to *cis*-regulatory elements (CREs), target genes and downstream regulatory networks. Variant-to-CRE identifies disease relevant CREs *in silico*, while CRE-to-gene and gene-to-network are experimentally inferred using TAP-seq (targeted Perturb-seq) and Perturb-seq. Lower part outlines the T cell transduction protocol used. Disease abbreviations: IBD, inflammatory bowel disease; UC, ulcerative colitis; AS, ankylosing spondylitis; RA, rheumatoid arthritis; SLE, systemic lupus erythematosus; MS, multiple sclerosis; PBCh, primary biliary cholangitis; PSC, primary sclerosing cholangitis; T1D, type 1 diabetes. (**B**) Genome-wide overview of the immune-disease variants (first track), CREs (second track), and promoters/genes (third track) tested in this study. Variants are shown with their GWAS association significance, CREs with ATAC-seq signal for naive CD4+ T cells 16 hours after activation with CD3/CD28^7^, and promoters/genes with expression level in naive CD4+ T cells in transcripts per million (TPM).

### Selecting immune-disease relevant CREs (variant-to-CRE)

We prioritized candidate CREs (cCREs) by intersecting immune disease variants with genomic regions displaying increased chromatin accessibility (ATAC-seq) upon TCR stimulation (CD3/CD28) compared to the naive state^7^. This yielded 844 disease-linked cCREs overlapping 4,724 GWAS variants across 14 immune and autoimmune traits, including inflammatory bowel disease (IBD), allergic disease, chronic liver disease, and systemic autoimmune disorders (IEU OpenGWAS) (Fig. 1; fig. S1A; table S2). We further added 181 cCREs that overlapped strong CD4+ T cell eQTL variants^8^ or gene regulatory networks (GRNs) ^22,23^, and 7 manually selected cCREs (materials and methods). This resulted in a total of 1,032 cCREs (Fig. 1B; table S3).

### TAP-seq links immune disease CREs to their downstream targets (CRE-to-gene)

TAP-seq provides higher sensitivity than Perturb-seq to detect the typically weak effects of enhancers and silencers, but requires an *a priori* list of candidate target genes for readout^15^. We selected candidate genes in a recall-oriented approach that prioritizes breadth over precision using five complementary prediction strategies: (i) proximity within 100 kb of the cCRE (3,703 links), (ii) the nearest up- and downstream genes to the cCRE if they fall outside of the 100 kb window (715 links), (iii) CD4+ T cell specific GRN connections^22,23^ (1,322 links), (iv) ENCODE-rE2G predictions^21^ (1,124 links; table S4), and (v) CD4+ T cell eQTL links^8^ (123 links) (Fig. 2A). This yielded 4,928 cCRE-gene pairs with an average of 5 links per cCRE. Many predicted links (34%) were supported by at least two approaches, and the resulting gene list was strongly enriched for T cell function (fig. S1B).

**Fig. 2:**
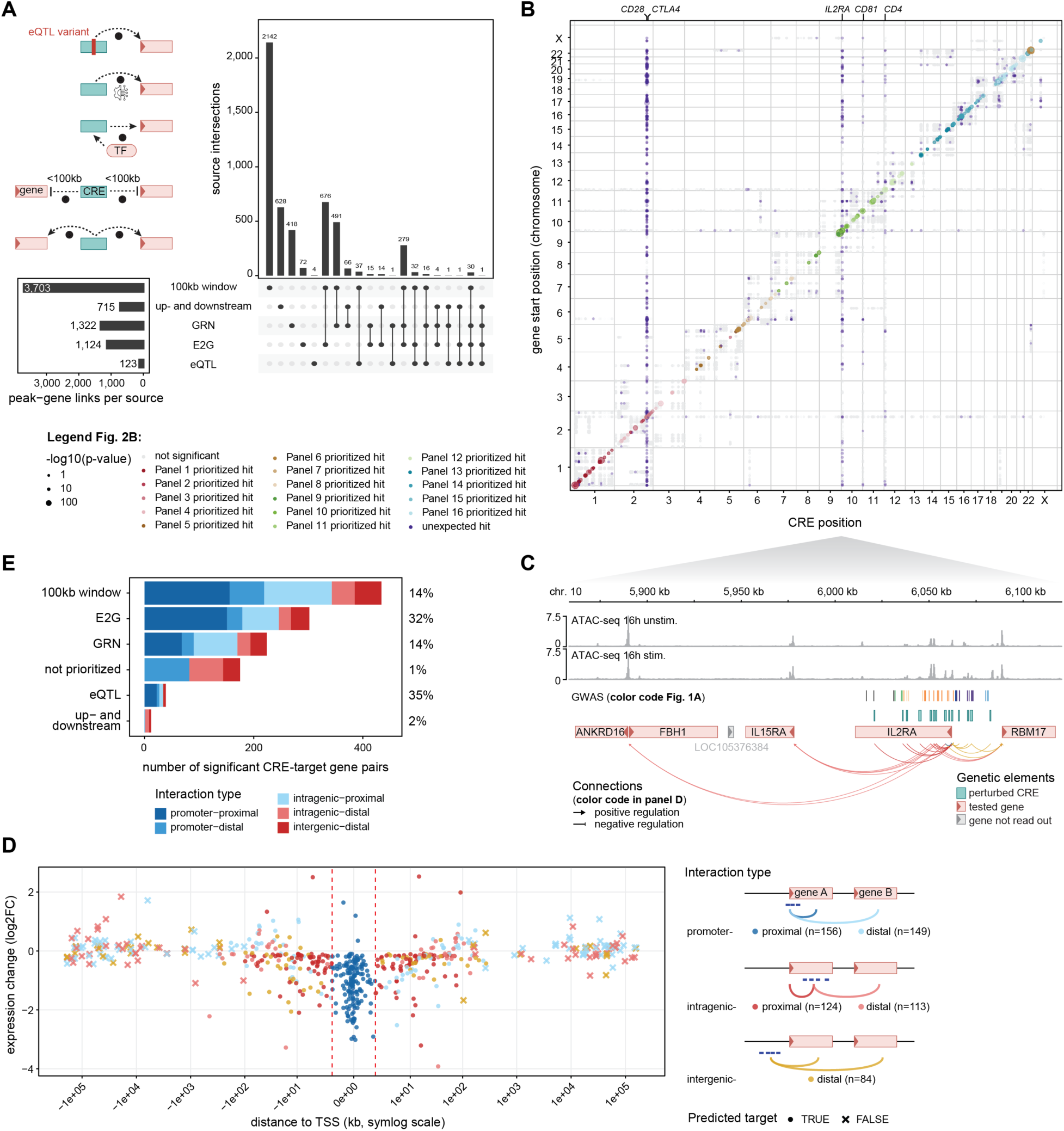
A large-scale targeted Perturb-seq (TAP-seq) screen identifies target genes of immune-disease relevant CREs. (**A**) Putative target genes of each immune-disease relevant CRE were predicted using five orthogonal approaches as described in the main text. The left panel shows a schematic of each of the five sources, the UpSet plot shows the intersection of the five sources, and the boxplot contains the absolute number of predictions from each source. (**B**) Genome-wide view of all identified associations from the CRE screen as identified by TAP-seq. The color code represents different experiments (pairs of gRNA libraries and TAP-seq target gene panels). Positive control promoters (*CD28*, *CTLA4*, *IL2RA*, *CD81*, *CD4*) were marked. (**C**) Example locus spanning a ∼250 kb region on chromosome 10 containing the *ANKRD16*, *FBH1*, *IL15RA*, *IL2RA*, and *RBM17* gene. The different tracks represent ATAC-seq data from different stimulation time points, as well as GWAS variants, CREs, and genes. Red and green arrows represent significant associations identified by TAP-seq and link the perturbed CRE with the gene that was read out with TAP-seq. TAP-seq connections with regions outside the locus are not shown. (**D**) Log fold change (LFC) and distance to TSS for significant TAP-seq CRE-gene links, colored by CRE interaction type t (promoter, intragenic, intergenic) and the relative position of the target (proximal, distal). The red dotted line represents a 1.5 kb region around the transcriptional start site. (**E**) Number of significant CRE-gene pairs identified by each of the five CRE-gene link prediction strategies. Percentages indicate the relative proportion of validated CRE-gene predictions for each prediction.

In the full CRE screen (design principles in supplementary text; figs. S1-3; table S5; table S6; table S7; table S8), we obtained 1,454,021 high-quality cells with at least one gRNA assigned (83% of all cells). This resulted in an average of 938 cells with uniquely assigned gRNA per cCRE (fig. S3A). Promoter perturbations, included as positive controls targeting genes at various expression levels (*CD28*, *CTLA4*, *IL2RA*, *CD81*, *CD4*), were robust across all 16 library-panel pairs (fig. S3F). After excluding 612 associations derived from control promoter perturbations and 200 associations across chromosomes (which are more likely to reflect *trans*-interactions), we identified 626 significant *cis*-regulatory interactions (Fig. 2B; table S9). We identified on average 1.6 target genes per CRE (27% upregulated, 73% downregulated upon CRISPRi perturbation) and the identified target genes showed on average 1.6 CRE regulating them. At the immune-relevant *IL2RA*/*CD25* locus, TAP-seq identified multiple variant-harboring CREs with blood and lymphoid eQTL support^18,19,24–26^ whose regulatory effects were consistent with long-range enhancer activity, a known *IL2RA* super-enhancer^10,27^, and additional events of positive and negative regulation (on *RBM17*, *IL15RA*, *ANKRD16*, *FBH1*) (Fig. 2C). An additional example is the *TNF* locus, where we identified multiple CREs regulating *TNF*, *LTA*, *LTB*, *LST1*, and *NCR3*, including an annotated enhancer-like element^28^, and a previously unknown silencer for *LTB* and *NCR3* (fig. S4).

### Targeted methods outperform distance as a strategy to predict CRE-gene links

We evaluated CRE-gene prediction strategies. *In silico*-predicted CRE-gene links were 13-fold more likely to be experimentally validated compared to non-predicted links (11% versus 0.9%; Fig. 2E). eQTL-based links showed the highest validation rate (35%) for few predicted links, while the 100 kb window yielded the largest absolute number of validated links (n = 434) but with low precision. ENCODE-rE2G predictions accounted for nearly 300 validated hits (32% validation rate), supporting substantial generalization beyond the K562 training data^21^.

To distinguish between different classes of CREs (including promoters, enhancers, and silencers) and disentangle them from CRE-independent effects, we grouped significant CRE-gene pairs by genomic context (promoter, intragenic, intergenic) and proximity to the nearest target gene transcription start site (TSS) (proximal, distal) (Fig. 2D; materials and methods for definitions). Most validated eQTL links were proximal (74%), whereas GRN-derived links, with a low overall validation (14%), were enriched for distal interactions (34%; fig. S5A). As most prediction strategies contain a distance component and significance thresholding cannot be unified across methods, these comparisons are confounded by the selection criteria. Still, all prediction strategies outperformed simple distance-based approaches (100 kb window, up- and downstream) in either precision or specificity for distal regulation, underscoring the value of prediction methods grounded in regulatory evidence for identification of distal CREs.

### Identification of enhancer-like regulatory interactions

To identify enhancer-like interactions (defined here as a CRE that up-(enhancer) or downregulates (silencer) the expression of a gene without acting through the promoter of that gene), we further examined the five interaction classes using functional annotations (chromatin accessibility, histone modifications, Hi-C data) (Fig. 3A). Promoter-proximal CREs behaved as canonical promoters (strong downregulation, promoter-like chromatin features), whereas intergenic-distal CREs showed hallmarks of long-range enhancer activity (H3K27acHi, H3K4me1Hi, H3K4me3Low). Within the remaining categories (promoter-distal, intragenic-proximal, intragenic-distal), enhancer regulation is not readily distinguishable from technical and biological confounders (indirect regulation through gene-gene interactions; transcriptional roadblocking via dCas9-binding in transcribed regions^29,30^; CRISPRi spreading^31^). To resolve these possibilities, we retained only distal effects occurring without any proximal effects that could mediate them. Within this refined set, we observed a distance dependent decrease in enhancer chromatin accessibility, H3K27ac and H3K4me1 signals, and TAP-seq fold-change, consistent with a decline in enhancer strength as distance increases (Fig. 3B; Fig. 2D). Enhancer-like CREs showed higher Hi-C contact frequencies with their target promoters than distance-matched non-significant CREs, were enriched among high-resolution promoter-capture Hi-C interactions^32^, and were typically located within the same topologically associating domain^33^ (Fig. 3C-D; fig. S5B). CREs located beyond 1 Mb from their target gene were indistinguishable from non-significant elements across chromatin and Hi-C features (Figs. 3B-C; fig. S5C), indicating limited direct *cis*-regulatory activity at such distances. This led us to a total of 106 high-confidence enhancer-like *cis*-regulatory interactions across 85 CREs and 89 target genes, comprising 84 activating (enhancer) interactions and 22 repressive (silencer) interactions.

**Fig. 3:**
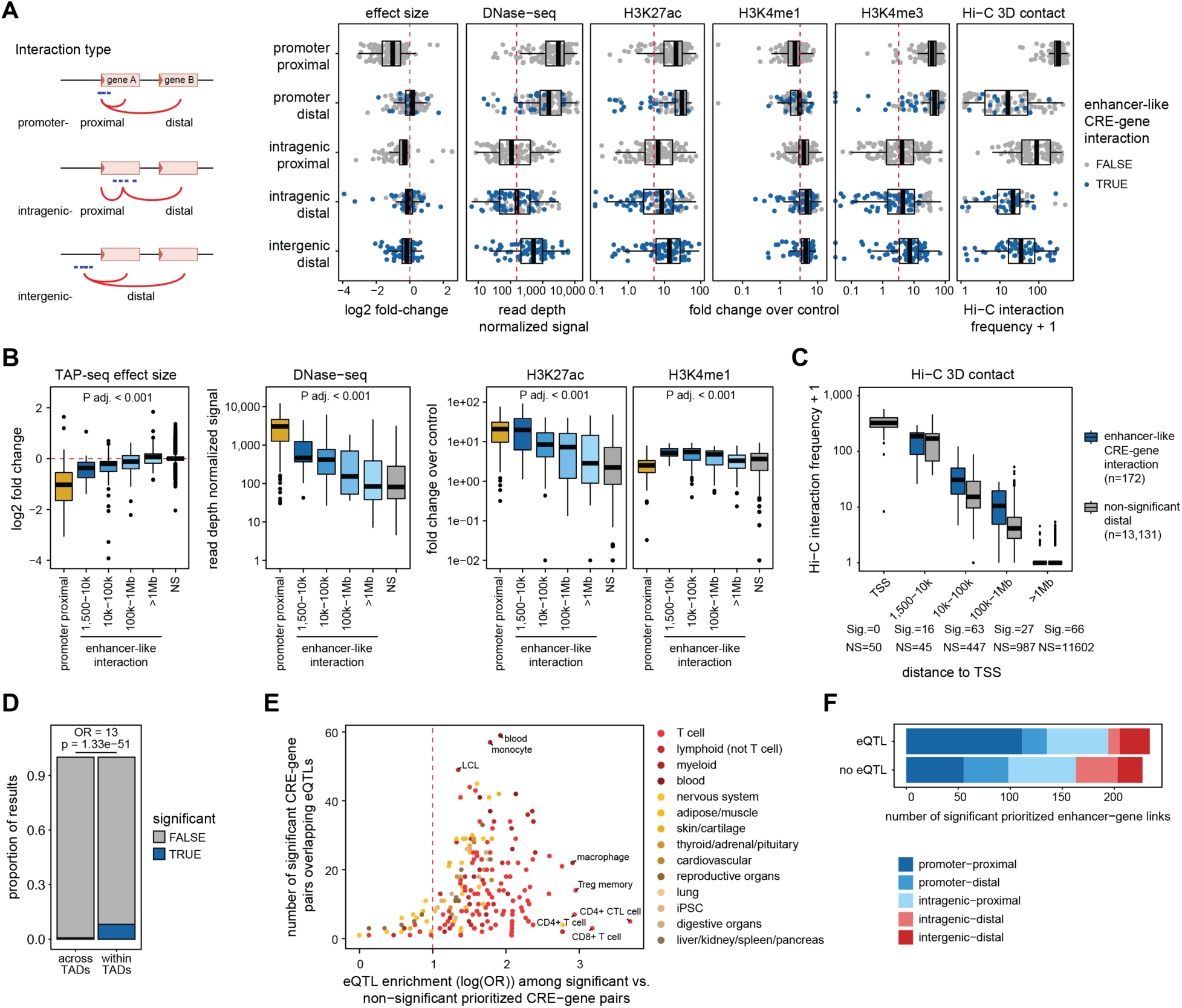
Identification of enhancer-like CRE-gene interactions by overlapping TAP-seq data with chromatin, Hi-C, and eQTL information. (**A**) Hits from the CRE screen were subdivided into five interaction classes based on genomic context of the perturbed CRE (promoter, intergenic, intragenic) and the relative position of the identified target gene (proximal, distal) (left, schematic representation). Right: TAP-seq effect size, DNase-seq, H3K27ac, H3K4me1, H3K4me3 and Hi-C 3D contact frequency for each identified TAP-seq hit, stratified by interaction class. Enhancer-like CRE-gene interactions (color code) were identified as described in the main text. (**B**) Stratification of effect sizes and chromatin properties by distance to transcriptional start site (TSS) for enhancer-like CRE-gene interactions and promoter-proximal interactions as reference. Distance is measured from the center of the perturbed CRE to the nearest annotated TSS of the respective gene and CRE-gene pairs are grouped into distance to TSS bins. If a CRE was linked to multiple genes, only the interaction with the closest TSS was retained. Number of CRE-gene pairs per interval: Promoter-proximal (n = 154); 1,500 - 10 kb window (n = 15); 10 kb - 100 kb window (n = 48); 100 kb-1 Mb (n = 22); >1 Mb (n = 50); not significant (NS, n = 173). (**C**) Relationship between Hi-C interaction frequency and distance to TSS for both significant enhancer-like CRE-gene interactions (sig.) and non-significant tested distal CRE-gene interactions (NS). Distance bins were defined as in panel B. Number of CRE-gene pairs per bin: Promoter-proximal (sig. = 0, NS = 50); 1,500-10 kb window (sig. = 16, NS = 45); 10 kb-100 kb window (sig. = 63, NS = 447); 100 kb-1 Mb (sig. = 27, NS = 987); >1 Mb (sig. = 66, NS = 11,602). (**D**) Proportion of enhancer-like CRE-gene links that are significant if the CRE and gene are located within the same topologically associated domain (TAD), versus if the CRE and gene have a TAD boundary in between them (across TADs). Odds ratio (OR) and p-value from Fisher’s exact test of enrichment. (**E**) eQTL enrichment comparing significant versus non-significant prioritized CRE-gene links. Plot shows log odds ratio (OR) of enrichment (x-axis) vs. absolute number of CRE-gene links supported by an eQTL for 280 datasets, colored by tissue. (**F**) Number of significant *a priori* prioritized hits with and without eQTL support across the five interaction types.

### Functional CRE-gene interactions are enriched for T cell-specific eQTLs

We assessed overlap between CRE-gene pairs with eQTLs from 281 eQTL datasets (table S10; Fig. 3E). The strongest enrichment of significant and prioritized CRE-gene links was found for CD4+ T cell-specific eQTLs (log odds ratio (OR) 3.7) ^34^, while the largest absolute overlap was with eQTLs from whole blood (n = 59) ^35^ (fig. S5D). More generally, blood eQTLs showed greater concordance with our CRE-gene observations than those from non-immune tissues (mean log(OR) 1.9 for blood cells and 1.8 for T cells, compared with 0.9 for internal organs and 1.2 for reproductive tissues) (Fig. 3E). More than half of the significant and prioritized CRE-gene links (51%, n = 234) overlapped with an eQTL, however, the majority of these corresponded to promoter-proximal effects (n = 111) (Fig. 3F). Overall, CRE–gene links are strongly enriched for immune-relevant regulatory variation.

### Genome-wide Perturb-seq in primary CD4+ T cells (gene-to-network)

To systematically map gene-to-network relationships, we performed genome-wide Perturb-seq targeting 7,664 genes, including all genes expressed in CD4+ T cells, transcription factors^36^, and target genes from the CRE screen (Fig. 1; screen design in Supplementary Text; fig. S6; table S11; table S12). Whole transcriptome libraries were sequenced to a mean of 13,679 UMIs per cell across 2,607,658 cells. After filtering, we retained 2,050,896 high-quality cells with assigned gRNAs (78% of cells with ≥1 gRNA; 99.8% of gRNAs detected in ≥10 cells), corresponding to a median of 134 cells per gRNA, or 402 per perturbation. Of uniquely assigned gRNAs, 79% showed significant on-target knockdown (Fig. 4A), consistent with another genome-wide Perturb-seq screen in primary CD4+ T cells (72%; ref. ^37^). With collapsed guides, knockdown efficiencies were comparable in primary CD4+ T cells (ours and ref. ^37^), below immortalized cell lines^38^, and above iPSCs^39^ (Fig. 4B).

**Fig. 4:**
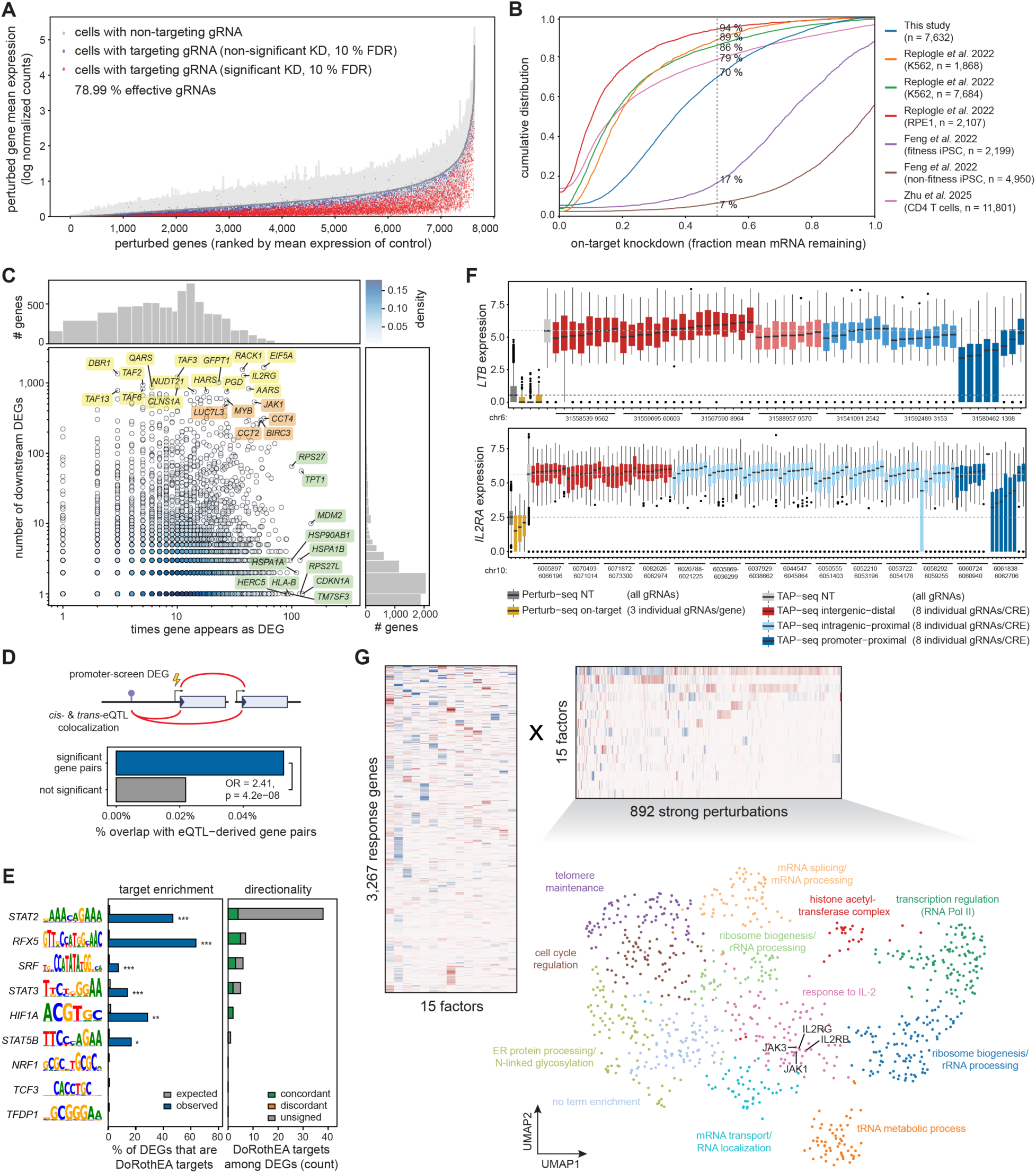
Genome-wide promoter Perturb-seq in primary CD4+ T cells. (**A**) On-target knockdown efficiency per gRNA, quantified by contrasting the mean log-normalized expression of the targeted gene between cells carrying that gRNA and non-targeting gRNAs (gRNAs achieving significant knockdown in red, non-significant knockdown in blue, and non-targeting controls (± s.d.) in gray). (**B**) Comparison of on-target knockdown efficiencies for different genome-wide promoter Perturb-seq screens in K562^38^, RPE1^38^, iPSCs^39^ and primary CD4+ T cells (this study and ref. ^37^), quantified by the ratio of mean log-normalized target gene expression in perturbed versus non-targeting control cells. (**C**) Gene regulatory connectivity in the promoter screen. Scatter plot showing the relationship between how often genes appear as differentially expressed genes (DEGs) across knockouts (x-axis) and the number of DEGs they cause when knocked out (y-axis). Points colored by density. Annotated genes highlight the top perturbations with broadest transcriptional impact (yellow), frequently dysregulated genes (green), and hub genes (orange, high in both dimensions). Log scales used for both axes; marginal histograms show distributions. (**D**) Schematic representation (top) and enrichment (bottom) of overlap between gene-gene pairs identified in the promoter screen and pairs derived from colocalized *cis*- and *trans*-eQTLs in blood. Significant target-response gene combinations from the promoter screen were more likely to overlap eQTL-derived pairs than non-significant pairs (OR = 2.41, P = 4×10-8). (**E**) Enrichment of DoRothEA transcription factor (TF) targets among promoter screen DEGs. Only TFs with ≥50 DoRothEA targets (confidence levels A-C) and ≥5 DEGs (adjusted P <0.1, |log2 fold change| >0.5) are shown. Left: Observed (blue) versus expected (gray) percentage of DEGs that are DoRothEA targets (one-sided Fisher’s exact test). Right: Concordant (green) if expression change matches DoRothEA regulatory direction, discordant (orange) if opposite, unsigned (gray) if no direction annotated in DoRothEA. *P <0.05; **P <0.01; ***P <0.001; n.s., not significant. (**F**) Comparison of perturbation effects between the CRE and promoter screens for two example genes (*LTB* and *IL2RA*). For each gene, expression is shown for non-targeting controls and individual gRNAs from the promoter screen (3 gRNAs/gene) and CRE screen (8 gRNAs/CRE), stratified by CRE interaction type. Horizontal dotted lines indicate the median of the CRE and promoter screen non-targeting gRNAs. (**G**) Embedding of 892 strong perturbations (>10 DEGs, Fig. S5F) using the factor model MOFA. Heatmaps show the two matrices learned by MOFA: Left, with the 3,267 response gene loadings across 15 factors; right, with scores for the 892 strong perturbations across 15 factors describing the contribution of each gene on these factors. Positive values shown in red and negative ones in blue. For visualization, the genes in both matrices were ordered based on hierarchical clustering and factors were sorted by decreasing variance explained. The resulting perturbation × factor matrix was used to cluster perturbations and visualized using UMAP, with points colored by cluster and annotated by functional enrichment.

We compared multiple differential expression strategies and selected DESeq2 on pseudobulks for its optimal tradeoff between calibration and discovery (supplementary text; fig. S6C-E). This identified 96,762 differentially expressed genes (DEGs; FDR 0.1) across 7,300 sufficiently covered perturbations and 8,001 variable genes, of which 81% had at least one DEG, and 12% >10 DEGs (“strong perturbations”) (fig. S6B; fig. S6C; fig. S6F; table S13). Perturbations with the highest number of downstream genes included TAF genes and *IL2RG*, reflective of their central roles in T cell functioning. Frequently dysregulated genes (heat shock proteins: *HSPA1B*, *HSP90AB1*, *HSPA1A*; and p53 pathway: *TM7SF3*, *MDM2*, *CDKN1A*) likely reflect general stress responses to perturbation (Fig. 4C). Effect sizes correlated well with bulk CRISPR data in CD4+ T cells^40^ (fig. S6D), and perturbation-response pairs were enriched among *cis-trans* blood eQTLs^41^ (Fig. 4D; table S14) and DoRothEA transcription factor targets^42^ (Fig. 4E; fig. S7). Comparison of effect sizes in the promoter- and CRE screen showed that promoter effects (i.e. promoter screen on-target and CRE screen promoter-proximal) result in the strongest downregulation, with TAP-seq giving clear advantage in sensitivity of detection (Fig. 4F).

We decomposed expression changes from 892 strong perturbations (fig. S6F) into latent factors representing co-regulated gene sets using MOFA (Fig. 4G; fig. S8A; table S15) ^43,44^. This revealed distinct transcriptional programs, including a JAK-STAT signaling-enriched factor driven by *STAT2*, *IRF9*, *IFNAR2*, *IL2RG*, and *JAK3* perturbation (fig. S8B-C). Clustering perturbations by factor loadings identified groups affecting related biological processes, such as transcriptional regulation, cell cycle control, and IL-2 response (driven by IL-2 receptor subunits *IL2RG*, *IL2RB*, *JAK1*, and *JAK3*) ^45^ (Fig. 4G).

### Propagating immune disease variant effects through regulatory networks (variant-to-CRE-to-gene-to-network)

We next constructed regulatory cascades from disease-associated variants by integrating the CRE and promoter screens. Of the 89 genes affected by high-confidence enhancer-like CREs, 41 (46%) showed DEGs when perturbed in the promoter screen, connecting to a total of 815 genes (effects downstream of CRE target genes in the promoter screen are defined as “hop 1”; materials and methods) (Fig. 5A). We further propagated the network with effects downstream of hop 1 genes (“hop 2”), leading to a highly interconnected immunoregulatory network encompassing 56% (4,356/7,815) of all tested genes. This integration facilitated the assignment of biological processes to CREs and diseases (fig. S9), demonstrating that disease-associated CREs connect to broad transcriptional programs through regulatory cascades (Fig. 5A).

**Fig. 5:**
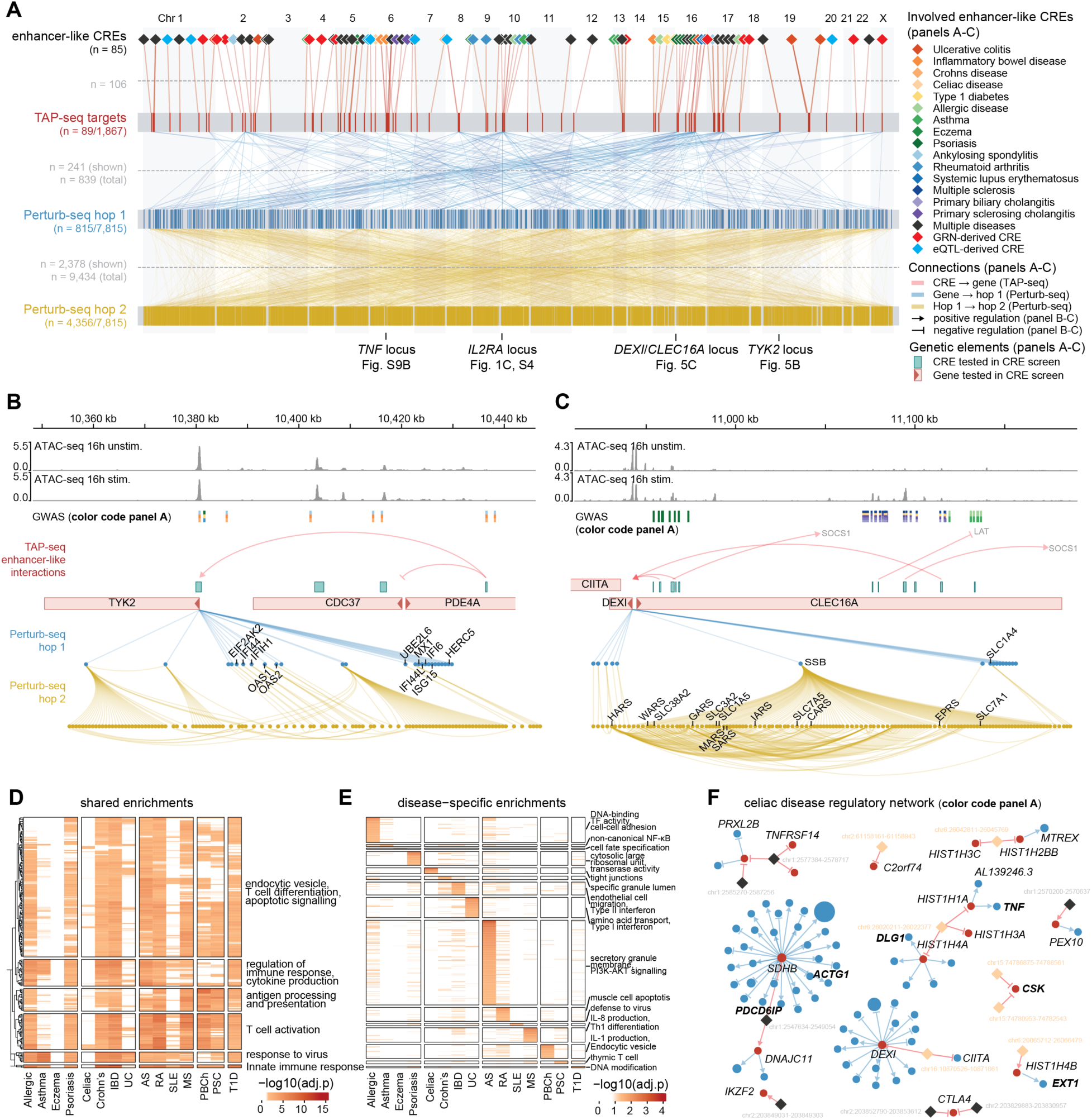
Integration of CRE- and promoter screen data for variant-to-CRE-to-gene-to-network. (**A**) Map of regulatory cascades as determined with TAP-seq and Perturb-seq. Genes and CREs are ordered by genomic position. Top row shows 85 enhancer-like CREs with significant target genes within 1 Mb, colored by associated disease trait. Second row shows TAP-seq tested genes (89 of 1,867 with enhancer-like hits; red, active; gray, inactive). The third and fourth rows show genes involved at Perturb-seq hop 1 (blue) and hop 2 (yellow). Lines between layers represent significant connections (adjusted P <0.1, |log2FC| ≥0.2, only connections to the top 20 targets per gene are shown). (**B**) Regulatory cascade at the *TYK2* locus. An enhancer-like CRE intronic to *PDE4A* regulates *TYK2*. *TYK2* perturbation in the promoter screen propagates to a canonical type I interferon-stimulated gene program (involved gene names indicated). Tracks show ATAC-seq, GWAS variants, and regulatory connections. Legend in panel A. (**C**) Regulatory cascade at the *DEXI*/*CLEC16A* locus. Three enhancer-like CREs associated with psoriasis and other immune diseases converge on *DEXI* without affecting *CLEC16A*. Network propagation identifies *SSB* as a downstream hub with enrichment for amino acid transport and tRNA aminoacyl-ligase activity (gene names indicated in lower part of the panel). Tracks and annotations as in panel B. (**D**) Heatmap of GO term enrichments (adjusted P <0.05) shared by >5 diseases, colored by -log10(adjusted p). Rows clustered by distance and split into main clusters, columns grouped by disease class, annotated based on representative GO terms. (**E**) Heatmap of GO term enrichments (adjusted P <0.05) unique to diseases, colored by -log10(adjusted p). Rows ordered by p-value, columns grouped by disease class, annotated based on representative GO terms. (**F**) Celiac disease network. Color code as shown in panel A. CREs associated with Celiac disease only (orange) or multiple diseases including celiac disease (black) are shown. CRE screen identified genes are indicated (black), as well as tight junction related (table S16) CRE screen and promoter screen hop 1 genes (black bold). Thresholds: adjusted P <0.1, |log2FC| ≥ 0.1.

One example of this is the *TYK2* (tyrosine kinase 2) locus (Fig. 5B). A CRE intronic to *PDE4A*, associated with IBD, ankylosing spondylitis, and Crohn’s disease, reduced *TYK2* (but not *PDE4A*) expression at 56 kb distance (LFC −0.07), identifying *TYK2* rather than the host gene as the likely regulatory target. Coding variants in *TYK2* confer protection against immune-mediated diseases^46^ and the protective allele of a Crohn’s disease variant reduces *TYK2* expression^8^. Our results provide functional evidence that a distal CRE can modulate disease risk through *TYK2* expression, suggesting an additional mechanism through which noncoding variants may act. *TYK2* is targeted by deucravacitinib, a selective inhibitor approved for psoriasis and suggested for use in other immune-mediated diseases including IBD^47,48^. Direct *TYK2* knockdown revealed a canonical type I interferon-stimulated gene program (including *MX1*, *OAS1*/*2*, *ISG15*, *IFI6*/*44*/*44L*/*H1*), consistent with *TYK2*’s role in *JAK*-*STAT* signaling. Together, our data place noncoding immune-disease variants upstream of *TYK2* and connect *TYK2* modulation to a downstream inflammatory pathway in activated CD4+ T cells.

The *DEXI*/*CLEC16A* locus exemplifies resolution of ambiguous GWAS target gene assignments (Fig. 5C). Variants within *CLEC16A* introns at 16p13 are associated with multiple sclerosis and other autoimmune disorders, yet act as eQTLs for neighboring *DEXI*, leaving the causal target unresolved^49^. Three enhancer-like CREs, overlapping with the *CLEC16A* intronic variants, converged on *DEXI* regulation while leaving *CLEC16A* unaffected, implicating *DEXI* as the target in CD4+ T cells. Network propagation revealed that *DEXI* knockdown altered SSB expression (hop 1), which regulated 115 downstream genes (hop 2; confirmed in ref. ^38^) enriched for amino acid transporters (including *SLC7A5*, *SLC1A5*, *SLC3A2*) and tRNA aminoacyl-ligases (including *WARS*, *HARS*, *EPRS*). These data link disease-associated noncoding variants to proliferative and metabolic programs via *DEXI* and *SSB*.

### Convergence of immune-disease variants on shared transcriptional programs

Although risk variants are distributed across the genome, they are thought to collectively contribute to disease by regulating gene programs. Using our variant-to-CRE-to-gene-to-network approach, we sought to identify convergent and disease-specific programs. For each disease, we aggregated implicated CREs and compiled genes connected through local (CRE-gene) and downstream (hop 1) regulatory effects. Gene Ontology analysis revealed substantial sharing of core immune programs (response to virus, T cell activation, and cytokine production were enriched in at least six diseases), consistent with the overlap of CREs across diseases and reflecting common immune regulatory architecture (Fig. 5D; table S16). In addition to shared programs, each disease network exhibited clear distinct enrichments (Fig. 5E; table S16). For example, genes downstream of multiple sclerosis-associated CREs were enriched for IL-8 production (adjusted P = 0.002), which is upregulated in serum and PBMCs from multiple sclerosis patients^50^. We found multiple celiac disease-associated CREs that converged on genes involved in tight junction related processes (including *DLG1*, *ACTG1*, *CSK*, *PDCD6IP*; top enrichment adjusted P = 1.4 × 10⁻⁷). This pathway is central to intestinal barrier dysfunction and is actively being explored as a therapeutic target^51^, illustrating how dispersed noncoding risk variants can converge on coherent disease-specific programs (Fig. 5F).

## Discussion

We present a genome-scale perturbation-based map of coding and noncoding genome function in primary human CD4+ T cells, a key cellular context for (auto)immune disease risk. By integrating *in silico* predictions with large-scale experimental testing, we established a framework that covers the entire chain from immune-disease variants to CREs, target genes, and downstream regulatory networks.

CRE-gene interactions were predominantly local, with modest effect sizes that required the sensitivity of TAP-seq. The impact of a CRE is not only shaped by its local effect, but also by the network downstream of its targeted genes. Network propagation using genome-wide Perturb-seq showed that the genes regulated by these CREs result in broad transcriptional responses that can reach 56% of all genes within two connections. Although immune-disease risk variants are distributed across many genomic loci, many affect shared downstream processes, including cytokine signaling and T cell activation. At the same time, we observed disease-specific programs, such as tight junction regulation in celiac disease, and combinations of these programs determine disease specificity.

Benchmarking CRE-gene linking strategies showed that ENCODE-rE2G and eQTLs had similar levels of agreement in CD4+ T cells and outperform simple distance-based approaches for identifying CRE-gene interactions, underscoring the value of computational inference. Despite being trained predominantly on K562 CRISPRi data, ENCODE-rE2G predictions showed general transferability across cell types while still requiring cell type-specific training for specific predictions. No single strategy outperformed the others in all metrics (precision vs. distal reach vs. absolute yield), highlighting the need for integrating orthogonal strategies.

Although Perturb-seq and TAP-seq enable systematic interrogation of regulatory elements, interpreting their results is inherently complex. By integrating 3D genome architecture and chromatin state information, we refine CRISPRi-based maps to identify high-confidence enhancer-like CREs, that cannot be reliably resolved using CRISPRi alone due to limited genomic resolution and the potential spreading of repressive effects. Through systematic comparisons of single-cell and pseudobulk analytical strategies, we showed that analytical choices can substantially influence the regulatory effects identified in Perturb-seq screens, highlighting the unmet need for standardized analytical frameworks. While our analysis focuses on activated CD4+ T cells, how these regulatory relationships propagate across T cell effector states will require future perturbation screens integrating coding and noncoding layers in those states.

By functionally mapping the coding and noncoding genome in the same cellular context, we deliver both a resource and a generalizable framework for dissecting disease mechanisms at scale. This variant-to-CRE-to-gene-to-network approach (i) can be transferred to other disease contexts where relevant model systems exist, (ii) provides training data for variant-to-function efforts, and (iii) suggests how to remove a blind spot in virtual cell models by extending their predictive value to noncoding genetic variation.

## Acknowledgments

We thank the EMBL Genomics Core Facility (GeneCore) for consultation and sequencing and Ralf Gilsbach (DZHK) for sequencer access. We thank the EMBL Flow Cytometry Core Facility for support and flow cytometry services. Vadir Lopez-Salmeron, Devon Jensen, and Cynthia Sakofsky (Waters Biosciences) for experimental advice and support. John Baeten, Andrew Boddicker, Ben Krajacich, Oliver Stephan, and Ewoud Ouwerkerk (Element Biosciences) for experimental advice and managerial support. Radu Raputeanu and Celia Alda Catalinas (GSK) for experimental and analysis advice. Tilmann Buerckstuemmer, Adam Krejci, Anke Loregger and Lukas Badertscher (Myllia Biotechnology) for experimental and analysis advice. Blagoje Soskic (Human Technopole), Carla Jones and Olivier Bakker (Sanger Institute) for data sharing and feedback on study design. Jonathan Pritchard, Alexander Marson, Emma Dann (UCSF, Stanford), and Brian Clarke (DKFZ) for discussions. Eugene Katsevich (UPenn) and Timothy Barry (Boston Children’s Hospital, Massachusetts General Hospital) for input on applying SCEPTRE. We thank all members of the lab of LMS for support and feedback.

## Funding

LMS was supported by grants from the Open Targets Consortium (project OTAR2063), by the National Human Genome Research Institute of the National Institutes of Health (RO1HG011664 & UM1HG011972), and the Dieter Schwarz Foundation Endowed Professorship. JME acknowledges support from the Applebaum Foundation, the BASE Research Initiative at the Betty Irene Moore Children’s Heart Center, and the NIH-NHGRI IGVF Consortium (UM1HG011972). ARG was supported by grants from the National Institutes of Health (R01HG011664 and UM1HG011972). AC received an EIPOD4 Fellowship from the EC Horizon2020 MSCA (847543).

## Author contributions

DPIM, AC, GT, DS, and LMS. conceptualized the project. DPIM, DS, and JY performed experiments. YM and MK generated gRNA libraries. AC, ARG, SS, DPIM, JB, CF, BR, and SZB performed bioinformatic analysis. DPIM, AC, ARG, SS, DS, and LMS wrote the manuscript. All authors read and commented on the manuscript.

## Competing interests

LMS is co-founder and shareholder of Sophia Genetics, Recombia Biosciences, and LevitasBio. MK is co-founder and shareholder of Vivlion. AA and RAA are employees at Waters Biosciences. DS and LMS received materials from Waters Biosciences and Element Biosciences that were used in this study. JME has received speaking honoraria from GSK plc, Roche Genentech, and Amgen. All other authors declare they have no competing interests.

## Data, code, and materials availability

All code and data will be made publicly available upon final publication.

## Supplementary Material

Materials and Methods

Supplementary Text

Figs. S1 to S9

Tables S1-S16

