## Supplementary Materials for "Genome-scale mapping of variant, enhancer and gene function in primary human CD4+ T cells"

Materials and Methods
Supplementary Text
Figs. S1 to S9
Tables S1-S16

#### Materials and Methods

##### T cell sourcing

Frozen healthy donor naive CD4+ T cells were obtained from AllCells (Discovery Life Sciences) as custom bulk isolations. They were isolated by negative selection from Leukopaks of three healthy, female donors and provided as isolated cryopreserved CD4+/CD45RA+ naive helper T cells (table S1). Donors were negatively tested for HIV, HBV and HCV.

##### Cell culture, maintenance and activation

Primary naive CD4+ T cells were cultured in RPMI 1640 (Gibco, cat. no. 52400-025) supplemented with 10% FBS Supreme (PanBiotech, cat. no. P30-3031), 1x penicillin-streptomycin (P/S, Gibco, cat. no. 15070063), 1 mM sodium pyruvate (NaPyr, Gibco, cat. no. 11360070), 0.1 mM non-essential amino acids (NEAA, Gibco, cat. no. 11140050), 55 µM 2-mercaptoethanol (Thermo, cat. no. 21985023), and 15 ng/µl recombinant human IL-2 (Peprotech, cat. no. 200-02). IL-2 was freshly added to the T cell medium. FBS was heat-inactivated at 56 °C for 30 min before use. One day post-thaw or plating, cells were activated using Human T-Activator CD3/CD28 Dynabeads (Gibco, cat. no. 11161) at a 1:1 cell-to-bead ratio and maintained at 0.5–2 x $10^6$ cells/ml.

LentiX HEK293T cells (Takara, cat. no. 632180) were cultured in DMEM (Gibco, cat. no. 41965-039) supplemented with 10% FBS Supreme, 2 mM GlutaMAX (Gibco, cat. no. 35050061), 1 mM NaPyr, 0.1 mM NEAA, and 1 x P/S. After thawing, cells were cultured for 48 h prior to medium change or passaging and maintained at 65–75% confluency.

##### Plasmids

*dCas9-KRAB constructs.* Plasmid pHR-UCOE-SFFV-dCas9-mCherry-ZIM3-KRAB was obtained from Addgene (plasmid #154473; a gift from M. Taipale). The open reading frames encoding ZNF264, ZNF324, ZNF554, or KRAB-MeCP2[52] were cloned into the pHR-UCOE-SFFV-dCas9-mCherry-ZIM3-KRAB backbone to replace the ZIM3-KRAB domain using Gibson assembly. A pHR-SFFV-dCas9-mCherry-KRAB construct containing the COX1 KRAB domain was generated by replacing BFP with mCherry in pHR-SFFV-dCas9-BFP-KRAB (Addgene plasmid #46911) using Gibson assembly.

*CROPseq-3Cs.* The Covalently Closed Circular synthesis (3Cs)[53] compatible CROPSeq-guide(F+E) vector was derived from CROPseq-Puro-F+E[15] by replacing the bacterial backbone of the CROPseq-F+E plasmid with that of pLentiCRISPRv2 (Addgene, #52961) using Gibson assembly. The spacer sequence between the U6 promoter and the SpCas9 scaffold v2 was subsequently replaced with the sequence 5'-GTT AAT TAA CCT TTT AAT TAA via Golden Gate cloning using Esp3I.

gRNA library cloning

*Pilot library.* The pilot gRNA library was generated for testing transduction protocols, experimental parameters and library preparation (fig. S2; table S8). Homology adapters were added to protospacer sequences (left: 5'-GAA GTG CCA TTC CGC CTG ACC TCG TCT CAC ACC, right: 5'-GTT TCG AGA CGA GGC TAG GTG GAG GCT CAG TG). Flanking regions contained BsmBI sites and Twist Universal Primer sequences. A 5′ G was added to all gRNAs. The gRNA pool was synthesized at Twist Bioscience, amplified using Twist universal forward (5'-GAA GTG CCA TTC CGC CTG ACC T) and reverse primers (5'-CAC TGA GCC TCC ACC TAG CCT) and cloned into CROP-seq (F+E)-Puro via BsmBI.

*CRE gRNA library.* The CRE gRNA libraries (fig. S1E; table S6) were synthesized as single-stranded DNA (oPool) at Integrated DNA Technologies. The libraries contained the following flanking sequences for 3CS cloning: left flanking sequence (5'-TGG AAA GGA CGA AAC ACC G) and right flanking sequence (5'-GTT TAA GAG CTA TGC TGG). The library was generated as a reverse complement for 3Cs cloning into CROPseq-3Cs.

*Promoter gRNA library.* The promoter gRNA library (Fig. S6B; table S11) was synthesized as single-stranded DNA at Twist Bioscience with flanking sequences compatible with 3Cs cloning. To improve cloning efficiency and library representation, the library was synthesized as a mixture in which 30% of gRNAs contained the standard left (5'-TGG AAA GGA CGA AAC ACC G) and right (5'-GTT TAA GAG CTA TGC TGG) flanking sequences (XX), 30% contained an additional

5′ G in the left flanking sequence (GX; 5′-GTG GAA AGG ACG AAA CAC CG), and 30% contained an additional 3′ A in the right flanking sequence (XA; 5′-GTT TAA GAG CTA TGC TGG A). The library was generated as a reverse complement for 3Cs cloning into CROPseq-3Cs.

*3Cs gRNA library cloning.* The libraries were constructed via hybridization and enzymatic conversion of a circular ssDNA template (CROPseq-3Cs), according to Wegner *et al.*[53] and Diehl *et al.*[54] except that the culture was grown at 30 °C for 18 h, and the final library plasmid DNA was isolated using a QIAgen Maxi Plus Prep kit. The CROPseq-3Cs ssDNA was produced following the protocol described by Wegner *et al.*[53] and Diehl *et al.*[54].

Lentivirus production

Lenti-X HEK293T cells were cultured to 65–75% confluency in T175 flasks using complete DMEM medium. Before transfection, the medium was replaced with 15 ml cOptiMEM (Gibco, cat. no. 51985-026, supplemented with 5% FCS, 0.1 mM NEAA, and 1× P/S). For transfection, 10 µg pMD2.G (Addgene, #12259) and 23 µg psPAX2 (Addgene, #12260) were combined with 32 µg CROPseq-3Cs plasmid with gRNA libraries or dCas9-KRAB plasmid. The DNA mix was combined with 112 µl P3000 in 4 ml OptiMEM, then mixed with 127 µl L3000 in 4 ml OptiMEM. The mixture was incubated at room temperature for 20 min and added dropwise to the cells. After 6 h, the transfection medium was replaced with complete DMEM containing 1:500 ViralBoost reagent. Viral supernatant was collected 48 h post-transfection, stored at 4 °C, and replaced with fresh medium. A second harvest was performed on day 3 post-transfection and pooled with the first. The combined supernatant was filtered through a 0.45 µm membrane and concentrated 100× using Lenti-X Concentrator (Takara, cat. no. 631232) according to the manufacturer’s instructions. Concentrated lentivirus was used immediately or stored at −80 °C.

T cell transduction

Naïve CD4+ T cells were thawed and cultured overnight at $1 \times 10^6$ cells per well in a 24-well plate. The next day, cells were activated and transduced by spinfection. Briefly, 87.5 µl lentivirus encoding either dCas9 or gRNA was added to $1\times10^6$ cells in 262.5 µl medium, yielding a total volume of 350 µl per well (25% v/v). T cells and virus were collected by centrifugation at 800 g for 60 min at 37 °C, and cells were returned to the incubator. After 4–6 h, 1.5 ml culture medium was added. Puromycin selection was performed with 2 µg/ml puromycin (Gibco, cat. no. A111380), and blasticidin selection with 20 µg/ml blasticidin (Gibco, cat. no. A1113903). Transduction timelines were tested as shown in Fig. S2A.

Fluorescence-activated cell sorting (FACS) experiments

Unstained cells were washed with FACS buffer (PBS + 2% BSA) and collected by centrifugation at 400 g for 5 min at room temperature. Cells were resuspended in FACS buffer at 10–20 × $10^6$ cells/ml and passed through a 35 µm cell strainer (Corning, cat. no. 352235). Cells were stained with DRAQ7 (Biostatus, cat. no. DR77524) for live/dead staining. Samples were kept on ice until further processing.

Flow cytometry was performed using a Waters Biosciences LSR Fortessa or Symphony A3 analyzer, FACS on a BD Aria Fusion II with a 100 µm nozzle. Sorted cells were collected in complete RPMI medium. Data was analyzed using FlowJo v10.9.0. Cells for analysis and sorting were gated for mCherry (dCas9-KRAB-ZIM3-mCherry) and DRAQ7 (live/dead staining).

scRNA-seq experiments

BD Rhapsody HT Xpress 3' gene expression profiling was performed with a targeted cell recovery of 45,000 to 55,000 cells per lane according to the manufacturer's instructions until the end of Exonuclease I treatment (Capture and cDNA Synthesis, protocol version 23-24252). From here, the TAP-seq and Perturb-seq libraries were processed differently (materials and methods 'TAP-seq library preparation' and 'Perturb-seq library preparation').

To store leftover beads for subsequent reactions ('bead recycling'), beads were resuspended in Bead Resuspension Buffer (Waters Biosciences), heat denatured at 95 °C for 5 min, followed by shaking incubation at 1,200 rpm for 10 s at room temperature. Beads were magnetized, supernatant was removed and the beads were resuspended in 200 µl cold Bead Resuspension Buffer (Waters Biosciences).

TAP-seq library preparation

TAP-seq libraries were prepared as described[15] with several modifications. Instead of using 10x Genomics 3′ scRNA-seq cDNA as input for PCR1, bead-bound cDNA from the BD Rhapsody HT Xpress system was used. To compensate for lower amplification efficiency in an on-bead PCR, the PCR1 volume was doubled from 100 µl to 200 µl. The PCR1 mix contained 8 µl 10 µM partial Read 1 primer, 5 µl 100 µM outer panel primer, 4 µl 12 µM outer CROP-seq primer, 83 µl $H_2O$, and 100 µL 2x KAPA HiFi HotStart ReadyMix (Roche, cat. no. KK2603). Rhapsody beads were magnetized, supernatant removed, and beads were resuspended in PCR1 mix. This mix was divided into four 50 µL reactions for the PCR run. PCR1 was run for 11 cycles to account for higher cell input as compared to the original method[15]. Reaction products were magnetized, and supernatant was purified using 1.5x AmpureXP (Beckman Coulter, cat. no. A63881) with three ethanol washes. During the final wash, all four reactions were combined and eluted in 50 µl Elution Buffer (Qiagen, cat. no. 19086). TAP-seq PCR2 was performed as described[15], using 10 ng of PCR1 product as input. PCR3 was modified to incorporate dual barcodes by replacing the Targeted 10x primer with custom i5-TruSeq Read1 primers. Final libraries were quantified using the Qubit HS dsDNA Assay (ThermoFisher, cat. no. Q33230). All primer sequences used for TAP-seq are listed in table S5 and table S7.

Perturb-seq library preparation

Perturb-seq libraries were generated in two steps: First, we prepared the CROP-seq gRNA library which is needed to assign gRNAs to cells; second, we prepared the whole transcriptome library. The CROP-seq gRNA library was prepared using the same nested PCR approach as for TAP-seq, leaving out the outer and inner panel primers in the first and second PCR respectively and incorporating an E-gel size selection (Thermo Fisher) post final cleanup to isolate the gRNA amplicon (~490 bp). The whole transcriptome libraries were prepared according to the manufacturer's instructions (BD Rhapsody HT Xpress mRNA whole transcriptome analysis (WTA), protocol version 23-24117). The gRNA library was added to the whole transcriptome library as a 2% spike-in before sequencing.

Sequencing of TAP- and Perturb-seq libraries

CRE and promoter screen libraries were sequenced using Element Biosciences AVITI. All libraries were first prepared as Illumina-ready libraries. Fragment sizes and concentrations were measured using an Agilent High Sensitivity DNA bioanalyzer and Qubit HS dsDNA assay. Subsequently, samples were pooled per chip and the pooled libraries were then converted to AVITI-ready libraries using the Element Adept Library Compatibility Kit v1.1 (Element Biosciences, cat. no. 830-00007). Promoter screen libraries were sequenced with a 1% PhiX spike-in, CRE screen libraries with a 5% PhiX spike-in. AVITI sequencing was performed using the Cloudbreak High 2x75 bp chemistry in High Throughput mode. The following sequencing cycles were applied for all libraries: 8 cycles [Index 1], 8 cycles [Index 2], 47 cycles [Read 1], 121 [Read 2]. CRE screen libraries were sequenced to an average depth of 7,000 reads per cell, promoter libraries were sequenced to an average of 27,000 reads per cell. Pilot experiments in Fig. S2 were sequenced using Illumina NextSeq 2000 without conversion.

CRE screen: GWAS selection

We compiled 17 GWAS datasets spanning 14 (auto)immune diseases[55-69] from the IEU OpenGWAS resource[70-72]. Only studies that included at least 10,000 individuals of European or mixed populations were included and for each trait, the study with the largest number of cases was selected. A list of included GWAS is provided in table S2.

CRE screen: CRE selection

CREs were selected based on their overlap with regions active in naive T cells and their association with autoimmune disease-related SNPs from GWAS. ATAC-seq data from naive, unstimulated, and CD3/CD28-stimulated (16 h and 5 d) CD4+ T cells [7] was then filtered for peaks that were more active in stimulated cells compared to non-stimulated cells. Peaks in the top 50% of most active peaks in at least one stimulated condition were retained. The resulting list of peaks were extended by 1 kb up- and downstream and analyzed for immune trait GWAS SNP enrichment using CHEERS[7], yielding a set of 844 disease-relevant CREs.

This list was expanded to include additional CREs that did not overlap with GWAS SNPs. CREs were added based on their overlap with enhancers in a CD4+ T cell-specific GRN[23]. The resulting CREs were filtered (peak-gene adjusted $P < 0.05$, peak-gene correlations $r > 0.67$ corresponding to

99.5th percentile of all GRN connections), adding 100 GRN-based CREs. Additionally, ATAC-seq and H3K27ac peaks overlapping with eQTLs[8] were included if the eQTL had a $p < 0.0005$, an absolute effect size >1, and a minor allele frequency >0.05. The related peaks were further required to show greater activity in stimulated cells over unstimulated cells. This added a further 81 eQTL-based CREs. Finally, we included two CREs from a GRN[23], and five CREs based on pilot data, bringing the total list to 1,032 unique CREs.

CRE screen: gRNA and gRNA library design

CRE-targeting gRNAs are difficult to design since gRNA design tools are optimized for promoter targeting gRNAs. We selected eight gRNAs per CRE using two different gRNA design tools (Fig. S1D): Four gRNAs were designed using CRISPRDesigner[73,74] within a 300 bp window around the ATAC-seq peak summit used for CRE selection[7]. If no summit was available, the 300 bp window was centered on the annotated CRE midpoint. Standard CRISPRDesigner filters (base composition, sequence complexity) were applied. gRNAs with <3 bp positional difference on the same strand were considered overlapping; in such pairs, the lower-scoring gRNA was removed. The top four non-overlapping gRNAs were retained. Four additional gRNAs were designed using CRISPick[75,76] within a maximum 2 kb region around the ATAC-seq peak, restricted to the annotated CRE boundaries. When no summit was present, the 2 kb window was centered on the CRE midpoint. CRISPick gRNAs were filtered using the same criteria applied to CRISPRDesigner, overlapping gRNAs (<3 bp difference on the same strand) were removed, and gRNAs overlapping selected CRISPRDesigner gRNAs were excluded. In cases where CRISPRDesigner or CRISPick were unable to design four gRNAs for a CRE, we supplemented gRNAs using the following strategy: If CRISPRDesigner produced fewer than four valid gRNAs, additional CRISPick gRNAs that passed our filters were added to reach eight gRNAs per CRE. In cases with still fewer than eight gRNAs per CRE, additional CRISPRDesigner gRNAs were added, including ones that initially were removed because of overlaps with other gRNAs (<3 bp difference on the same strand). After applying these rules, only 18 CREs had fewer than eight gRNAs designed.

Control promoters and CREs from the pilot study were targeted with four to eight gRNAs each. These control CRE-targeting gRNAs were taken from[14] or designed using guidescan[77], promoter control gRNAs from Dolcetto Set A[76]. In total, the CRE libraries contained 8,175 gRNAs targeting CREs, 30 non-targeting gRNAs[76], 34 CRE-targeting gRNAs from the pilot study, and 20 gRNAs targeting positive control promoters, resulting in 8,259 gRNAs. We additionally included 17 gRNAs targeting other manually selected loci which we didn't include in our analysis (in total 8,276 gRNAs). The CRE-targeting gRNAs were subdivided into 16 sublibraries, where each panel contained the 30 non-targeting gRNAs and 20 control promoter gRNAs; panel 1 additionally contained the 34 CRE-targeting gRNAs from the pilot study.

CRE screen: Target gene selection

Using the list of CREs (materials and methods 'CRE screen: CRE selection'), potential CRE target genes for TAP-seq readout were selected according to the following five selection strategies and were required to be expressed in at least five cells based on whole transcriptome data: (i) 100 kb window: All genes within 100 kb up- and downstream of the selected CREs were included. (ii) Up- and downstream: The nearest gene up- and downstream of each selected CREs was included, independent of the distance. (iii) GRN: Genes were incorporated based on regulatory network evidence from two CD4+ T cell-specific GRNs. From the activated CD4+ T cell network[23], gene-CRE links meeting an adjusted P <0.3 were retained when the corresponding CRE was part of the screened set. From a naive CD4+ T cell GRN[22], associations were included if they satisfied an adjusted P <0.3 and a peak-gene distance <250 kb, provided the linked CRE was selected for perturbation. (iv) eQTL: Genes were added on the basis of genetic evidence when a significant (adjusted P <0.05) naive CD4+ T cell eQTL signal[8] colocalized with a screened CRE. (v) ENCODE-rE2G: target genes were incorporated from ENCODE-rE2G enhancer-gene predictions[21] derived from 12 naive and stimulated CD4+ T cell biosamples (table S4), when predicted enhancer coordinates coincided with CREs selected for perturbation.

Applying these criteria yielded a list of 1,866 selected target genes for TAP-seq read-out. Additionally, a set of 107 genes relevant to T cell biology was cherry-picked (based on a gene list generated for a related T cell perturbation project by our collaboration partner Gosia Trynka, Sanger Institute) and added. Ten housekeeping genes were also included, chosen based on medium expression levels, low expression variability, distance from CRE targets (>2MB), and lack of perturbation effects in Replogle *et al.*[38].

Selected CREs and corresponding target genes were subdivided into 16 perturbation panels. Excluding controls and promoter targets, CREs were clustered according to shared target genes using pairwise Jaccard similarity (threshold >0.05). Clusters were subsequently distributed based on genomic position, with manual refinement to balance aggregate expression levels across panels (fig. S1C). The 107 manually curated genes, lacking CRE assignments, were allocated by genomic proximity to the nearest CRE-containing panel. Control CREs were placed in gRNA panel 16 (table S6), whereas promoter controls and housekeeping genes were represented in all panels.

CRE screen: TAP-seq primer panel design

Target-specific primers were designed using the TAP-seq primer design pipeline (v1.15.2)[15], implemented as an R package on Bioconductor. Inner primers were positioned 150–300 bp upstream of annotated polyA sites, and outer primers 300–500 bp upstream, based on polyA annotations derived from whole-transcriptome pilot data. Because transcript 3′ ends were occasionally misassigned by the pipeline, primer pairs were manually curated by visual inspection of scRNA-seq read coverage and reference annotations in IGV. Primer sets falling outside the predominant 3′ end (highest coverage) were replaced with alternative candidates from the TAP-seq output.

Primers were synthesized as individual oligonucleotides (IDT). Outer and inner primers were pooled equimolarly into separate panels. The CROP-seq vector amplification primer was incorporated during PCR1 and PCR2 setup.

CRE screen: Data processing and quality controls

Reads were processed with the BD Rhapsody v2.2 pipeline. For each donor and perturbation panel, sequencing lanes were merged and analyzed jointly, with estimated cell numbers from the BD Rhapsody scanner. Alignment was performed against a custom hg38 reference generated using make_rhap_reference_2.2.cwl, incorporating gene annotations and gRNA sequences. Expression matrices were log-normalized using default Seurat parameters. Doublets were identified with scDblFinder (v1.12.0)[78], with the expected doublet rate set according to scanner-based measurements, and removed prior to downstream analyses (fig. S3B).

CRE screen: gRNA assignment

To generate gRNA assignments for the CRE screen, we used SCEPTRE's mixture method[79]. This approach models gRNA UMI counts using a Poisson GLM to assign guides to cells, while accounting for cell-specific covariates such as sequencing depth and batch to control for ambient background contamination.

CRE screen: Perturbation identification

Seurat objects generated by the BD Rhapsody pipeline were analyzed for differential expression using SCEPTRE (v0.10.0)[80,81] within a Snakemake workflow. Analyses were performed per panel, and donor-specific results were combined. Default SCEPTRE settings were applied, ensuring robust calibration and false discovery rate control. Non-targeting cells served as controls, and two-sided tests were performed. Treatment and control groups were required to contain at least seven nonzero cells. Cells were further filtered to retain UMI and gene counts within the 0.01–0.99 percentile range. Analyses were conducted in low-MOI mode without imposing an additional distance threshold, and covariates (donor, panel, lane, chip) were included in the model. Significant target-response associations were defined using Benjamini-Hochberg correction at an FDR threshold of 0.1.

CRE screen: Interaction types and enhancer-like CRE-gene links

To generate a list of enhancer-like CRE-gene interactions, all significant interactions of intergenic CREs with target genes on the same chromosome (intergenic-distal) were selected. This list was supplemented with other significant distal interactions (intragenic-distal, promoter-distal) without evidence of an effect mediated through the CRE host gene. If an intragenic or promoter-overlapping CRE had an effect on the host gene, then a gene-gene effect was considered more likely as the mechanism behind the observed effect on a distal gene and the pair was not included as an enhancer-like CRE-gene interaction.

CRE screen: Chromatin enrichment analyses

All candidate enhancer gene pairs were annotated for chromatin activity assays and 3D contact measurements in T cells using data available on the ENCODE portal[82]. BigWig files for DNase-seq accessibility (ENCFF600IHC), and H3K27ac (ENCFF151LJA), H3K4me1 (ENCFF075XMI) and H3K4me3 (ENCFF546IYU) ChIP-seq were used to quantify the chromatin assay signal at each candidate enhancer. Signals per assay were summed up for each candidate enhancer and normalized by enhancer size for ChIP-seq assays.

High resolution Hi-C data for CD4+ T cells was downloaded in .hic format (ENCFF355VJW). Hic-straw v1.3.1[83] was used to extract SCALE normalized observed contact frequencies at 5000 bp resolution for each candidate enhancer gene pair. For a given pair, the contact frequencies between the Hi-C bins overlapping the center of the candidate enhancer and gene TSS were used to quantify 3D contact.

For TAD-enrichment analysis, BED files containing TAD boundaries for naive CD4+ T cells stimulated with CD3/CD28 were obtained directly from the authors of Onrust-van-Schoonhoven *et al.*[33]. The resolution for the Hi-C experiments was 50kb, so the boundary elements are 50 kb wide. CREs and target genes were classified into their respective TADs using GenomicRanges v1.56.1[84] and elements on TAD boundaries were removed. Enrichment of tested and significant interactions within and across TADs was calculated and significance was evaluated using a $\chi$-squared Test.

CD4+ T cell specific promoter-capture Hi-C (pcHi-C) data restricted with DpnII was obtained directly from the authors of Malycheva *et al.*[32]. pcHi-C data was provided at the level of the individual DpnII fragments (median fragment size 266 bp) and combined into 5 kb windows. We combined both and tested whether enhancer-like CRE-gene links that overlapped pcHi-C contacts were more likely to be significant using a Fisher exact test, testing only genes whose promoters had been included in the pcHi-C data.

<u>CRE screen: eQTL enrichment analyses</u>

We tested if there was an enrichment of eQTLs among the significant CRE-gene links in the CRE screen using blood-specific *cis*-eQTLs finemapped using SuSie (eQTLGen phase 2)[41] and 280 smaller datasets available on eQTL Catalogue[18] on 30th of July 2025 (table S10). Since CD4+ T cell-specific eQTL information was used in the prioritization of possible target genes, we calculated enrichment of eQTLs among significant prioritized hits compared to non-significant prioritized hits using the log odds ratio (OR).

Promoter screen: Gene selection

First, we selected all expressed genes from a single-cell RNA-seq dataset of activated (CD3/CD28) naive CD4+ T cells that we generated as part of this study. Cells with particularly high and low UMIs were removed, as well as cells with >20% mitochondrial reads. Raw count data was normalized with SCTransform and TPM normalized. We excluded all genes detected in <6 cells, with tpm <10, as well as non-protein-coding genes. We then added back all genes (coding or noncoding) from our list of predicted CRE targets, as well as all transcription factors annotated in the Hocomoco database v12[36] with TPM >1.

Promoter screen: gRNA design

For genome-wide promoter Perturb-seq, we used 3 independent single-targeting gRNA constructs, instead of a single dual-targeting construct as done previously[38]. Three gRNAs were chosen per gene from the following sources in decreasing order of preference: Dolcetto set A[76], Dolcetto set B[76], CRISPRi v2.1[85]. If less than 3 gRNAs were available from these three sources, we designed new gRNAs using CRISPick. If CRISPick couldn't find gRNAs, the gene was dropped. This resulted in 7,664 genes for which 3 gRNAs were designed. 5% of the library size was added as non-targeting control gRNAs from Dolcetto setA and setB. Dolcetto and CRISPick gRNA protospacers were 20 nt long, CRISPRi 2.1 protospacers were 19 nt long.

Promoter screen: Data processing and quality controls

Promoter screen sequencing data was processed using version 2.2 of the BD Rhapsody pipeline. Reads were aligned against a custom reference genome constructed using the `makeRhapReference` function of the BD Rhapsody pipeline, which included the full genome (hg38), gene annotations, and gRNA sequences. The expected cell numbers per donor were calculated based on BD Rhapsody scanner data and we extracted data for a total of 2,607,658 cells from 64 samples (donor, chip, panel combinations). All output Seurat objects containing scRNA-seq UMI counts per sample were combined into one Seurat object using memory-efficient on disk storage using the BPCells (v. 0.3.0), Seurat (v 3.5.0) and SeuratObject (v 5.1.0) packages for R (v. 4.4.1). For analyses in python, all scRNA-seq data was exported as an AnnData object using the `write_matrix_anndata_hdf5` function from the BPCells R package.

For analyses in R (SCEPTRE, limma), data was further processed and quality controlled using Seurat (v 5.3.0) and BPCells (v 0.3.1). Data were normalized with log1p before sketching a subset of 100,000 cells using `SketchData` with method `LeverageScore` to determine QC thresholds. We kept cells with >1,000 genes, <15% ribosomal reads and <10% mitochondrial genes.

Promoter screen: gRNA assignment

gRNAs were assigned to cells based on UMI counts using the SCEPTRE Nextflow pipeline using SCEPTRE (v. 0.10.3)[79-81] and ondisc (v. 1.2.0) for memory-efficient on disk data storage for R (v. 4.4.1). To import the scRNA-seq data into SCEPTRE, UMI counts for 2,393,341 cells passing QC thresholds (described in previous section) were exported as .mtx files with accompanying feature and barcode metadata files following 10x cell ranger standards to track cell barcodes, gene ids and names, and distinguish between gene and gRNA features. SCEPTRE's `import_data_from_cellranger` function was then used to create a SCEPTRE object with high-MOI settings from these files. gRNA assignment was performed using the mixture method by executing the SCEPTRE Nextflow pipeline until the `assign_grnas` step. A custom model formula that included the following covariates was used: `~log(response_n_umis) + log(response_n_nonzero) + log(grna_n_umis + 1) + log(grna_n_nonzero + 1) + chip + panel + percent_rb + response_p_mito + CellCycle_G2M + early + ICOS_CD38 + Mito + CTLA4_CD38 + Poor_Quality + iTreg_5d4h + Th1_5d`. Default values were used for other SCEPTRE parameters.

gRNA assignment identified 1,410,422 cells with one uniquely assigned gRNA, 640,474 cells with multiple gRNAs, and 342,445 cells with no assigned gRNAs. For downstream analysis, we retained only cells with unique gRNA assignments (1,410,422 cells; 59%).

The gene set used for downstream analyses comprised all promoter-targeting perturbation genes together with additional highly variable genes identified using Scanpy's `highly_variable_genes` function (v. 1.11.5). Highly variable genes were selected independently within each sequencing batch based on normalized and log-transformed expression values from all recorded cells. Mitochondrial genes were excluded, resulting in a final set of 8,001 genes retained for downstream analysis.

<u>Promoter screen: gRNA and on-target knockdown efficiency</u>

To assess knockdown efficiency at the individual gRNA level, the mean log-normalized expression of each target gene was compared between cells carrying a given targeting gRNA and cells carrying non-targeting control gRNAs, following the analysis of Zhu *et al.*[37]. For each targeting gRNA, a Welch's t-test was performed against the non-targeting control population, and resulting p-values were adjusted using the Benjamini-Hochberg procedure. Overall, 79% of gRNAs exhibited a significant on-target knockdown, defined by an adjusted $P < 0.1$ and a negative t-statistic (Fig. S5B).

Knockdown efficiency was further evaluated at the target level using the fraction of remaining mRNA, defined as the ratio of mean log-normalized target gene expression in perturbed cells to that in non-targeting control cells (Fig. 4a). This metric was used to compare on-target knockdown efficiency across existing CRISPRi perturbation datasets, featuring K562 and RPE1 cell lines[38], iPSC[39], and primary T cells[37] (Fig. 4B).

<u>Promoter screen: Differential testing</u>

Differential expression (DE) testing was performed using DESeq2[86], accessed via the pertpy Python package[87], following the analysis of Zhu *et al.*[37]. Single-cell expression profiles were aggregated into pseudobulk samples by summing mRNA counts from cells sharing the same donor and gRNA identity, yielding up to nine biological replicates per target (three gRNAs per target across three donors). Pseudobulks with fewer than five cells as well as perturbations with less than 3 biological replicates were excluded from the analysis, resulting in a total of 7300 tested perturbations.

DESeq2 modeled gene-level counts using a negative binomial generalized linear model with a design formula that included the number of cells per pseudobulk (log10-transformed), donor identity, perturbation identity, and the fraction of counts mapping to ribosomal protein genes, resulting in the following design formula: `~ log10_n_cells + donor + target + ribosomal_percentage`. Perturbation targets were partitioned into batches comprising 50 perturbations together with all non-targeting control pseudobulks, and DESeq2 models were fitted independently for each batch[37]. For each perturbation, $\log_2$ fold-change estimates were obtained from the fitted models, and statistical significance was assessed using Wald tests contrasting targeting with non-targeting pseudobulks. Resulting p-values were adjusted for multiple testing using the Benjamini-Hochberg procedure.

Statistical calibration and false positive rates of DESeq2 were evaluated using a null experiment in which random sets of three non-targeting control pseudobulks were assigned as a pseudo-target group and contrasted against the remaining non-targeting pseudobulks. This design mirrored the discovery analysis, yielding pseudo-target groups with up to nine pseudobulks across donors and gRNAs, and uses the same cutoffs and design formula. We evaluated 300 random sets of NT gRNAs, and the resulting p-value distributions closely followed the null hypothesis (fig. S5E). At an adjusted $p < 0.1$, an overall false discovery rate of 0.03% was observed (fig. S5C).

<u>Promoter screen: Other tools</u>

In addition to DESeq2, multiple differential testing approaches were evaluated, including commonly used tools such as SCEPTRE[79-81] and limma[88], as well as Wilcoxon rank-sum tests applied to both single-cell and pseudobulked data. These complementary methods were used to assess the robustness of differential expression results to differences in statistical modeling assumptions, data aggregation strategies, and control of false positive rates.

*SCEPTRE*. Differential expression analysis was performed using the SCEPTRE Nextflow pipeline in the high-MOI setting and the same model formula as for gRNA assignment (materials and methods 'Promoter screen: gRNA assignment'). To jointly analyze gRNAs targeting the same promoter, `grna_integration_strategy` was set to "union." To construct negative control targets for running calibration checks, `calibration_group_size` was set to 3 to match the 3 targeting gRNAs per promoter, and 200,000 calibration checks were performed by setting n_calibration_pairs accordingly. For comparative analyses, the calibration results were filtered to include only genes considered in the other 2 differential expression approaches and randomly downsampled to 100,000 pairs. Power checks were performed using the gene associated with each targeted promoter as positive controls. Differential expression tests for the full screen were performed using the massive *trans* analysis setting, which tested every promoter perturbation for effects on every gene. Resulting p-values were corrected using the Benjamini-Hochberg procedure and an adjusted p <0.1 was used to identify significant effects.

*Wilcoxon single-cell and pseudobulk*. As a baseline, Wilcoxon rank-sum tests were performed using Scanpy's `rank_genes_groups` function (v. 1.11.5)[89] at both the single-cell and pseudobulk levels on log-normalized expression data. For single-cell analyses, all perturbations represented by more than five cells were tested. Calibration was assessed using a one-versus-rest analysis of non-targeting control gRNAs and revealed substantial p-value inflation, consistent with pseudoreplication from treating individual cells as independent observations (fig. S5C). As an alternative to single-cell testing, a pseudobulk Wilcoxon approach was evaluated. Pseudobulk profiles were normalized and log-transformed prior to testing, and the same pseudobulk construction and filtering criteria as used for DESeq2 were applied, but without covariate adjustment.

*Limma*. We used R package limma (v 3.60.3) to run differential expression at single cell level. Cells with 1 gRNA assigned were retained for analysis, and non-targeting gRNAs with at least 15 significant DEGs in the Wilcoxon single-cell analysis were removed. To enable analysis of the large dataset, cells were divided into 5 splits based on perturbation, where cells with non-targeting gRNAs were added to each split. We subsetted the counts matrix to the 8,001 variable genes. We normalized to counts per million based on size factors and differential expression was run with the following design matrix: `gene + sequencing_run + nr_UMIs + percent.mt + percent.rb + CellCycle.G2M + early + ICOS.CD38 + Mito + CTLA4.CD38 + Poor.Quality + iTreg_5d4h`. We used `eBayes(fit, trend = TRUE, robust = TRUE)` to extract DEGs for each perturbation and used the adjusted P <0.1 as significance threshold.

Promoter screen: overlap with eQTL-derived *cis-trans* gene pairs

To validate the promoter screen DEGs, we tested if significant (FDR <0.1) target-DEG pairs were enriched for gene pairs derived from eQTL mapping. The eQTLGen Consortium constructed gene pairs where the same genetic signal affects a local gene (*cis*-eQTL) and a distal gene (*trans*-eQTL) based on formal colocalization[41]. We filtered the eQTL pairs for the set of 8,001 variable genes and performed a Fisher exact test to calculate the enrichment of significant compared to non-significant promoter screen pairs.

Promoter screen: Transcription factor analysis

To assess whether transcription factor (TF) perturbations preferentially affect their known target genes, we performed enrichment analysis using the DoRothEA database[42]. TF-target interactions were obtained from OmniPath[90], retaining interactions with DoRothEA confidence levels A, B, or C. For each perturbed gene in the Perturb-seq dataset, differentially expressed genes (DEGs) were identified using thresholds of adjusted P <0.1 and |log2 fold change| >0.5. Self-hits (where the perturbed gene itself is differentially expressed) were excluded. Genes annotated as TFs in the DoRothEA database were considered for enrichment analysis if they had at least 5 DEGs in Perturb-seq and at least 50 DoRothEA targets (levels A-C) among the 8,001 genes measured per perturbation. Enrichment of DoRothEA targets among DEGs was tested using a one-sided Fisher's exact test against the background of all measured genes. Directional concordance was evaluated for TF-target pairs with annotated regulatory direction in DoRothEA. A target was scored as concordant if an activated target was downregulated after knockout (log2FC <−0.5) or a repressed target was upregulated (log2FC >0.5). The opposite direction was scored as discordant. Targets without directional annotation were classified as unsigned. Statistical significance of directional concordance was assessed using a one-sided binomial test against the null hypothesis of 50% concordance among directionally annotated targets.

Promoter screen: gRNA-level knockdown comparison with CRE screen

To look into gRNA-level effects from both screens on the same gene, we selected 2 genes (*LTB* and *IL2RA*) that were targeted by several classes of CRE and significantly downregulated in the promoter screen. We selected cells with 1 assigned gRNA from either screen, normalized the counts with log1p and plotted the expression level in all cells carrying non-targeting gRNAs compared to the cells carrying individual targeting gRNAs.

Promoter screen: MOFA analysis

To identify perturbations with similar downstream effects, we fitted a Multi-Omic Factor Analysis (MOFA) model implemented in the MOFAflex python package (v. 0.1.0)[43,44] to the log fold changes obtained from DESeq2. We selected 892 strong perturbations (>10 DEGs with an adjusted P <0.1) and 3,267 downstream genes that were significantly affected by at least 10 perturbations and retained only log fold changes with an adjusted p <0.2 while remaining values were set to NA, ensuring that the factor model decomposition focuses on significant log fold changes.

To select an optimal number of factors, MOFA was run with 10 different random seeds using a Normal likelihood and varying the number of factors (5, 10, 15, 20, 25, and 30). To induce sparsity, a horseshoe prior was used for both the `weight_prior` and `factor_prior`. Training parameters were set to a learning rate of 0.005, `early_stopper_patience` = 1000, and a maximum of 20,000 epochs. The best performing model was chosen based on total variance explained.

The strong perturbations were subsequently clustered based on their low-dimensional embedding from MOFA (the factor matrix Z) using the Leiden clustering algorithm as implemented in scanpy (flavor = “igraph”) (v. 1.11.5)[89]. For visualization of the embedding and the resulting clusters, UMAP was applied using scanpy and annotated using GO enrichment (see below).

For each latent factor, functional enrichment analysis was performed on both perturbations and downstream genes using the top 10% of genes with the highest absolute weights. Enrichment was performed using Enrichr via the gseapy Python package (v. 1.1.11)[91] against curated gene set collections, including Hallmark, Reactome, KEGG, and Gene Ontology Biological Process annotations for humans. For each analysis, the background set consisted of all perturbations or all downstream genes included in the MOFA model, respectively. Enrichment results were filtered to retain terms with an adjusted $p < 0.05$ after multiple testing correction.

Promoter screen: Propagating genome wide signaling cascades

*CRE and promoter screen network.* We constructed a hierarchical network in which nodes represent enhancer-like CREs and genes, and edges represent regulatory interactions identified by the CRE screen or promoter screen. The network consists of four layers ordered by chromosomal position: CREs (layer 0), TAP-seq tested genes (layer 1), and two successive layers of variably expressed genes reached through downstream (hop 1 and hop 2) Perturb-seq interactions. Hop 1 includes all genes that are significantly (FDR <0.1) perturbed by the promoter perturbation. Hop 2, includes genes significantly differentially expressed (FDR <0.1; absolute log fold change >0.2) downstream of hop 1 genes. CRE-to-gene edges were drawn for significant enhancer-like interactions within 1 Mb of the target TSS. Gene-to-gene edges were drawn for Perturb-seq hits passing 10% FDR and $|\log2 \text{ fold-change}| \geq 0.2$, excluding self-perturbations. For visualization in Fig. 5A, edges per source gene were capped at the 20 most significant targets; genes in the subsequent layer were still activated based on the full uncapped set.

*Disease regulatory network (Fig. 5F).* CRE-gene interactions were filtered for significant, non-control *cis*-interactions resulting from CREs selected for association with a specific trait. Genome-wide promoter Perturb-seq differential expression results were filtered for an adjusted $p \leq 0.1$ and absolute log2fold change of $\geq 0.1$. Directed networks were constructed by linking (i) CREs to their primary target genes identified in the CRE screen and (ii) primary target genes to downstream differentially expressed genes identified in the promoter screen (hop 1). Self-loops and duplicate edges were removed. Node degree in hop 2 was calculated to estimate downstream connectivity, determining node size. Graphs were visualized using force-directed layouts implemented in R (igraph, tidygraph, ggraph). Edge color indicated direction of regulation.

Promoter screen: Cytopus enrichment analysis

For each CRE with significant enhancer-like TAP-seq interactions, a gene cascade was constructed by iteratively expanding through Perturb-seq gene-gene links: direct CRE-gene targets, first-hop Perturb-seq targets of CRE-gene links, and second-hop targets. At each cumulative depth, gene sets were tested for overrepresentation in Cytopus v1.3 immune cell-type gene set library (254 terms)[92] using one-sided hypergeometric tests with Benjamini-Hochberg correction (adjusted P <0.05). To evaluate TAP-seq specificity, a parallel cascade was built using all CRE-gene predictions in place of TAP-seq hits. To evaluate Perturb-seq specificity, two null models were applied at each depth: (1) replacing cascade-added genes with size-matched random draws from the variable gene background, and (2) preserving cascade topology but substituting random Perturb-seq source genes at each hop. Both nulls were run for 100 permutations per CRE.

Promoter screen: Gene Ontology enrichment analysis

We used the R package clusterProfiler (v 4.12.0) and the function `enrichGO()` to test for enrichment of gene sets for GO terms (Biological Process, Cellular Component, Molecular Function). Gene symbols were converted to Entrez IDs using the package org.Hs.eg.db (v 3.19.1). The set of 8,001 variable genes were given as background for all analyses. P-values were corrected for multiple testing with the Benjamini-Hochberg method and the reporting p-value cut-off was 1 (i.e. all enrichments given as output). Enrichments are reported sorted by adjusted p-value.

*Disease enrichments*. To identify disease-specific GO enrichments, we combined the genes downstream of the CREs linked to a disease. For each disease, significant (FDR <0.1) CRE-gene links were included (no filter for chromosomal distance) and linked to promoter screen results if they were significant (FDR <0.1). The set of promoter screen response genes was used as input for enrichment analysis, removing gene sets with 3 or fewer genes. GO enrichments for all diseases were combined and classified as 'shared' if they were significant (adjusted p <0.05) in more than 5 diseases, and 'specific' if they were significant in exactly one disease. Heatmap diseases were grouped manually. The shared heatmap was clustered with the `cluster_rows` and `cluster_row_slices` arguments of R package ComplexHeatmap (v 2.20.0). The disease-specific heatmap was ordered by maximum enrichment p-value.

*Perturbation enrichments*. Similar to disease-specific enrichments, we also ran enrichments for gene cascades from individual perturbations for all hop 1 and hop 2 genes downstream of one promoter perturbation at a time.

*MOFA enrichments*. To identify shared functionality of the perturbations that clustered together based on their loadings of the MOFA factors, we performed GO enrichment for each of the 12 perturbation clusters. Each gene set consists of the perturbations in each cluster. The UMAP visualization was annotated with the top-enriched (by adjusted p-value) GO term.

#### Supplementary Text

CD4+ T cell perturbation and activation protocol

To achieve efficient CRISPRi and library coverage in primary CD4+ T cells, we optimized delivery of the CRISPRi components (fig. S2A) and selected the most effective CRISPRi KRAB domain (fig. S2B).

*T cell transduction protocol.* Previous studies have established transduction strategies for pooled CRISPR screening in primary human CD4 T cells[93,94]. We compared these and modified versions thereof that varied in the timing of transductions, selection steps and activation cycles (fig. S2A). The strategies were evaluated based on cell yield, viability, reproducibility, and fraction of dCas9 expressing cells. Of the four strategies, a modification version of the protocol described in ref. [94] outperformed the others. In the optimized approach, CD3/CD28-activated CD4+ T cells were transduced sequentially with lentiviral vectors encoding dCas9-KRAB-mCherry and gRNAs expressed from a CROP-seq vector[15,95]. gRNA expressing cells were selected for by puromycin. After transduction and selection, cells were reactivated and analyzed via flow cytometry for viability and expression of dCas9 prior to single-cell RNA-seq analysis.

*CRISPRi system.* We targeted the promoter of the surface receptor CD4 with six dCas9-KRAB or dCas9-KRAB-MeCP2 versions[52], and read out CD4 protein using flow cytometry (fig. S2B). dCas9 fused to the ZIM3 KRAB domain led to the strongest reduction of CD4 protein levels (reduction of 81% relative to control gRNA containing samples) and was therefore used for all experiments.

We further determined optimal experimental parameters for TAP-seq in a pilot screen with regards to CRISPR knockdown efficiency, cell coverage, and read coverage (fig. S2C-G).

CRE screen design principles

Candidate *cis*-regulatory elements (cCREs) and target genes were grouped into 16 pairs of gRNA libraries and TAP-seq primer panels based on genomic proximity and gene expression levels (figs. S1C-E; table S5; materials and methods). Each cCRE was targeted with eight gRNAs (control promoters and manually selected cCREs with four gRNAs) (figs. S2E-F; table S6); combined with 30 non-targeting gRNAs, this yielded a final library of 8,259 gRNAs (fig. S2G; materials and methods).

Promoter screen design principles

The experimental parameters for the Perturb-seq promoter screen were identical to the TAP-seq CRE screen, using the same donors, CRISPRi machinery, and time points for transduction, activation, and single-cell readout (Fig. 1A). Three gRNAs were designed per gene (one gRNA per construct) and delivered as a single library comprising 23,983 gRNAs, including positive and negative controls (fig. S6A; table S11).

Promoter screen differential gene expression testing

Since best practices for differential gene expression analysis from genome-wide Perturb-seq do not yet exist, we systematically compared single-cell approaches (testing perturbation effects at the individual cell level) and pseudobulk approaches (aggregating counts across cells sharing the same gRNA and donor) using four differential expression methods: DESeq2[86], limma-trend[96], SCEPTRE[81], and Wilcoxon rank-sum tests (fig. S6C; materials and methods). Single-cell Wilcoxon tests showed substantial p-value inflation consistent with pseudoreplication from treating individual cells as independent observations (fig. S6C). Covariate-adjusted discovery methods such as limma-trend and SCEPTRE partially mitigated this issue on single cell data, yielding more on-target discoveries while reducing the total number of differentially expressed genes, consistent with a lower false-positive rate. However, calibration tests for these approaches remained inflated. In contrast, pseudobulk Wilcoxon tests effectively controlled false positives but were underpowered, resulting in markedly fewer discoveries and could not detect on target effects (fig. S6C). Pseudobulks Wilcoxon tests controlled false positives but were underpowered, yielding markedly fewer discoveries. DESeq2 applied to pseudobulk profiles[37] provided the best tradeoff between model calibration, on-target hit discovery, and total number of significant associations (fig. S6C). All methods showed concordant effect size estimates with bulk CRISPR knockout data in CD4+ T cells[40] (fig. S6D), though DESeq2 pseudobulk achieved the highest per-perturbation correlations. Based on these results, DESeq2 on pseudobulk profiles was used for all downstream analyses reported in the main text.

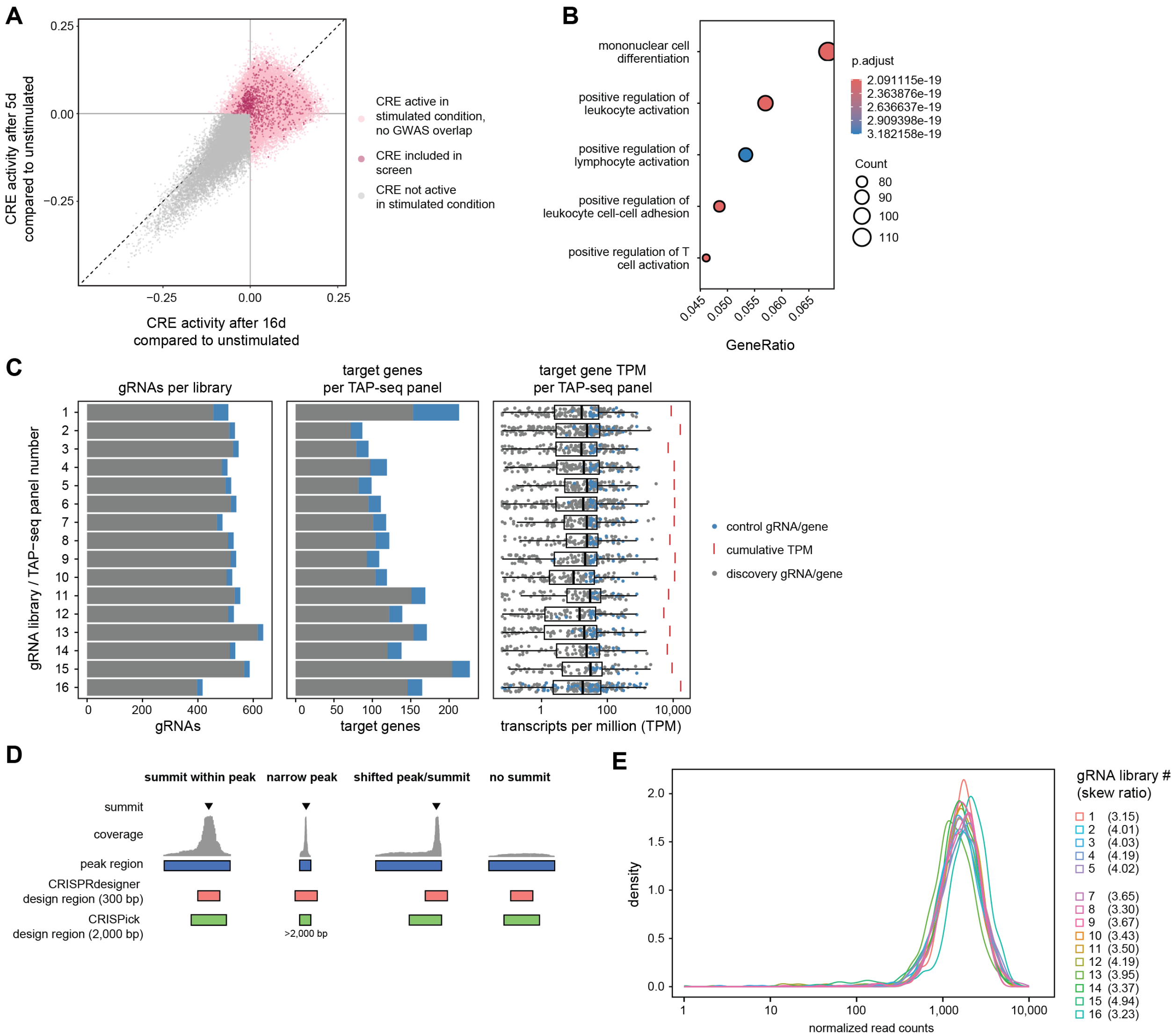


**Fig. S1: Design of the CRE screen gRNA libraries and TAP-seq target gene panels.** (**A**) CREs are shown based on increased activity after 5 or 16 d of stimulation relative to the unstimulated state. Only CREs showing increased activation in at least one stimulated condition were considered. From this set, CREs overlapping immune-disease GWAS variants were included in the screen and are indicated. (**B**) Top-5 gene ontology (GO) term enrichments in the CRE screen candidate target gene list as prioritized using the selection strategies shown in Fig. 2A. (**C**) The CRE screen gRNA libraries and TAP-seq target gene panels were subdivided into 16 sub-libraries or sub-panels (see materials and methods). From left to right: number of gRNAs per library, number of target genes per TAP-seq panel, and (cumulative) expression level in each TAP-seq panel. One library/panel pair (number 17) is not shown since it contains only control gRNAs/targets (non-targeting gRNAs, control enhancers, control promoters). TPM, transcripts per million. (**D**) Design strategy for CRE-targeting gRNAs used in this study was based on two publicly available tools, CRISPick and CRISPRdesigner (see materials and methods). (**E**) Normalized gRNA count distribution of each CRE screen gRNA sub-library. Skew ratio statistics for each sub-library are listed.

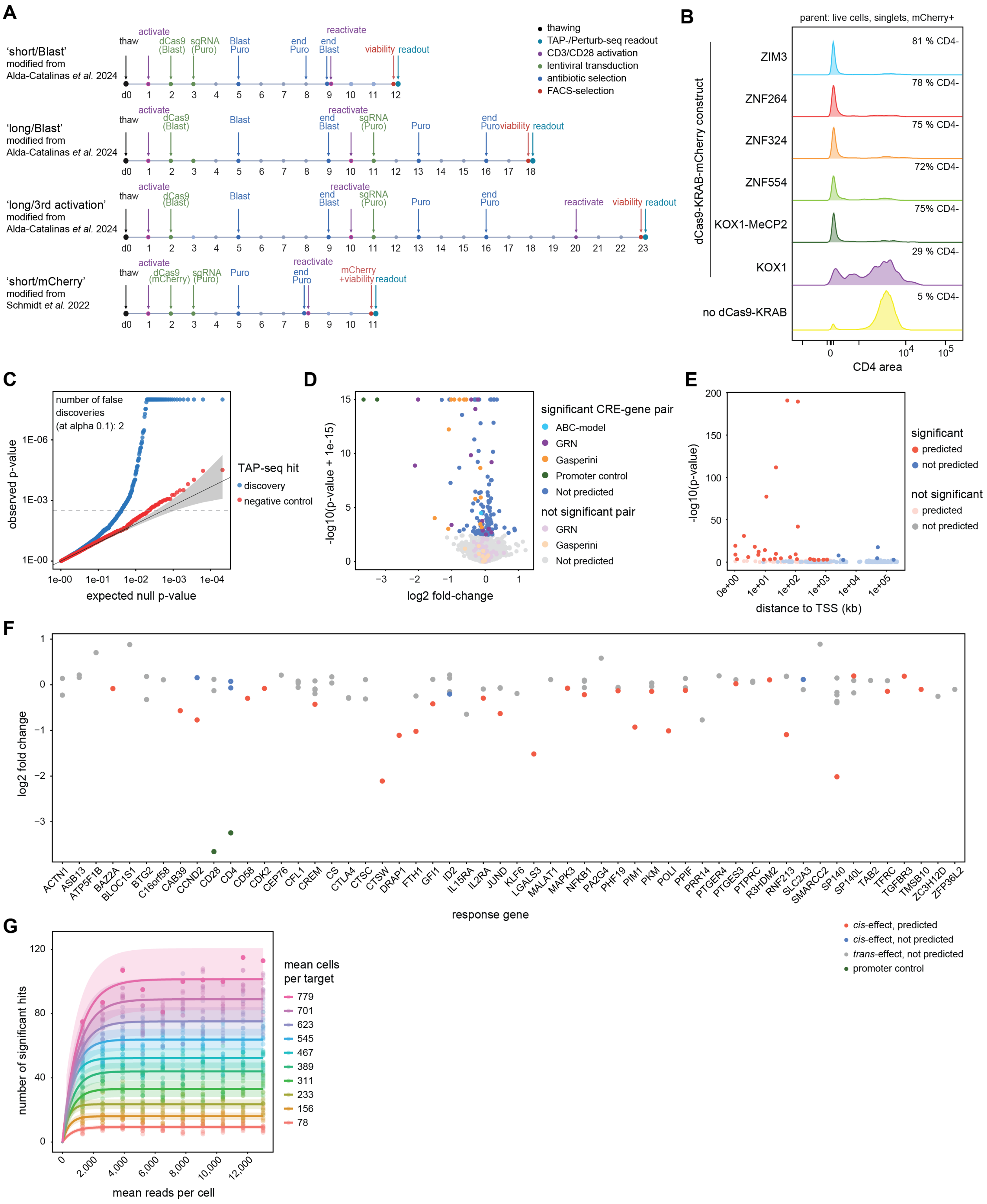


**Fig. S2: TAP-seq pilot studies define optimal design parameters of the large-scale CRE screen in CD4+ T cells.** (**A**) Schematic of the T cell transduction protocols tested in this study. Protocols were adapted from the indicated references[93,94]. Timelines show thawing (black), dCas9 and gRNA lentiviral transduction (green), T cell activation (purple), selection (blue), FACS enrichment (red), and readout (cyan). The 'short/mCherry' protocol[94] was used for both the CRE and promoter screens with slight adaptations (see materials and methods). (**B**) Comparison of dCas9 fusion proteins for CRISPRi in primary CD4+ T cells. CD4-targeting gRNAs or a non-targeting control gRNA were co-transduced with the indicated dCas9 constructs, and CD4 expression was assessed by FACS. Shown are the percentages of CD4-negative cells. (**C**) Expected null effect p-values plotted against observed p-values for negative controls and discovery hits. (**D**) CRE target gene effect size and scaled p-values, with CRE-target gene pairs colored according to their source and significance level. (**E**) Significance of CRE–gene associations as a function of genomic distance between each CRE and the transcription start site (TSS) of the regulated gene. (**F**) Effect sizes of enhancer perturbations on the indicated genes. CRE–gene pairs are colored to indicate whether they act in *cis* (same chromosome) and whether the effect was predicted *a priori*. (**G**) Downsampling analysis of pilot data using SCEPTRE. The dataset was downsampled to defined fractions of the full data, with 10 iterations performed per fraction. A negative exponential growth curve was fitted across fractions.

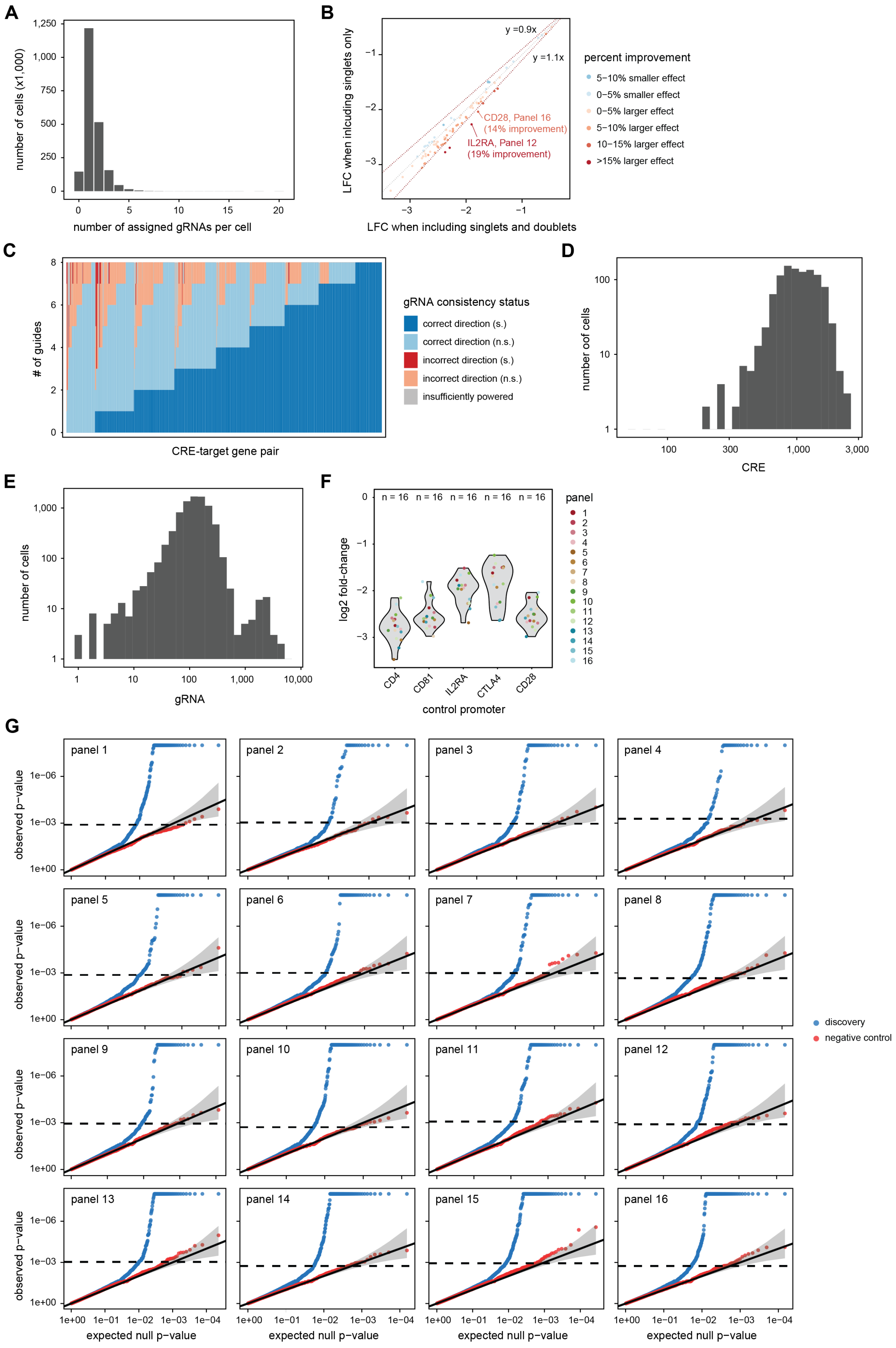

A
number of cells (x1,000)
number of assigned gRNAs per cell
B
LFC when inlcuding singlets only
LFC when including singlets and doublets
y =0.9x
y =1.1x
CD28, Panel 16 (14% improvement)
IL2RA, Panel 12 (19% improvement)
percent improvement
5−10% smaller effect
0−5% smaller effect
0−5% larger effect
5−10% larger effect
10−15% larger effect
>15% larger effect
C
### of guides
CRE-target gene pair
gRNA consistency status
correct direction (s.)
correct direction (n.s.)
incorrect direction (s.)
incorrect direction (n.s.)
insufficiently powered
D
number oof cells
CRE
E
number of cells
gRNA
F
log2 fold-change
control promoter
CD4
CD81
IL2RA
CTLA4
CD28
panel
G
observed p−value
expected null p−value
discovery
negative control

**Fig. S3: Additional results and quality metrics of the CRE screen.** (**A**) Number of gRNAs per cell according to the negative binomial model from SCEPTRE for the entire CRE screen. 50% of cells were assigned to one gRNA, 28% to multiple gRNAs, and for 22% of cells, no gRNA could be assigned. (**B**) Effect of including or excluding cell doublets as identified with scDblFinder[78] on measured log2 fold changes (LFC). Removing doublets increased on-target effects by 3.89%, with 56.6% of effects becoming stronger after removing doublets. (**C**) Per gRNA results from the CRE screen. For each identified CRE-gene pair, the numbers of gRNAs are shown which show a consistent or inconsistent direction (s, significant; ns, not significant; calculated using a two-sided t-test of the counts of a given gene for all cells assigned to a specific gRNA vs. all control cells). Of the significant ETPs, 93.1% were supported by at least 6 gRNAs (i.e. at least 6 gRNAs all showed a down-regulating effect, or at least 6 gRNAs showed an up-regulating effect). (**D**, **E**) Number of cells per CRE (F) or gRNA (G) for the full CRE screen. (**F**) Log2 fold-changes for each of the five positive control promoters (*CD4*, *CD81*, *IL2RA*, *CTLA4*, *CD28*) and for each of the 16 library/panel pairs. (**G**) Quantile-quantile (QQ) plots comparing the SCEPTRE calibration check and discovery analysis p-values against the expected null p-value distribution. Plotted on a negative log-transformed scale with gray regions corresponding to the 95% confidence bands.

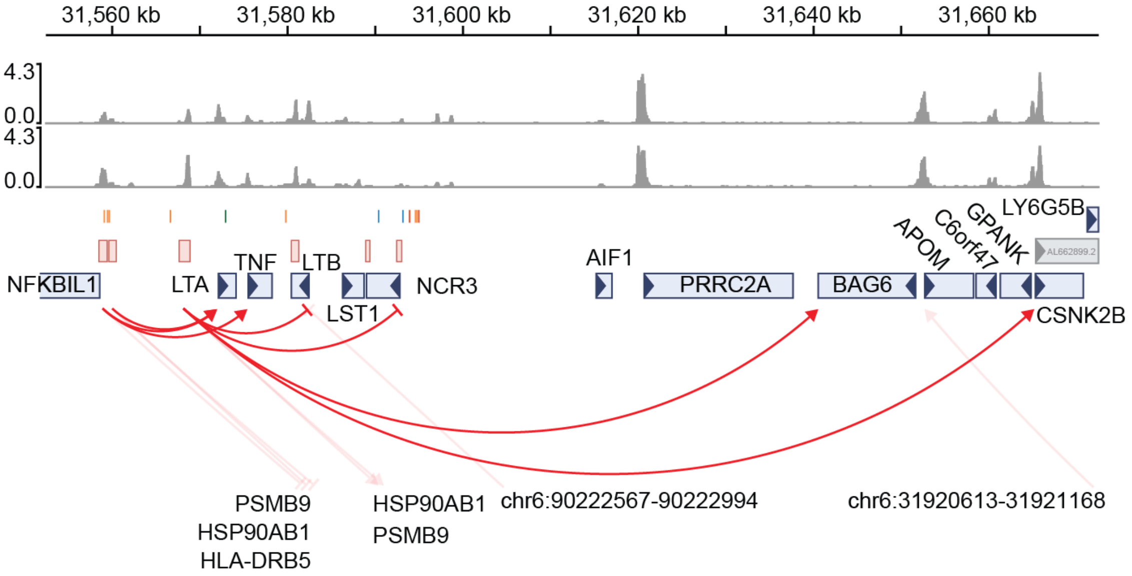


**Fig. S4: CRE-gene interactions at the *TNF* locus.** Genomic region spanning ~100 kb around the *TNF* locus on chromosome 6. The region contains *NFKBIL1*, *LTA*, *TNF*, *LTB*, *LST1*, *NCR3*, *AIF1*, *PRRC2A*, *BAG6*, *APOM*, *C6orf47*, *GPANK*, *CSNK2B*, and *LYG5B*, several of which are members of the *TNF* superfamily or are located within the extended MHC locus. Tracks show ATAC-seq signal across stimulation time points, immune-disease GWAS variants, prioritized CREs, and annotated genes. Significant TAP-seq associations are indicated by connecting lines; arrowheads denote positive associations and blunt-ended lines denote negative associations. Only enhancer-like CREs are shown. A previously annotated super-enhancer (chr6:31,567,298–31,589,539) is located in this region[28]. In addition to enhancer-like interactions, a novel silencer-like element regulating targeting *LST1* and *NCR3* was identified.

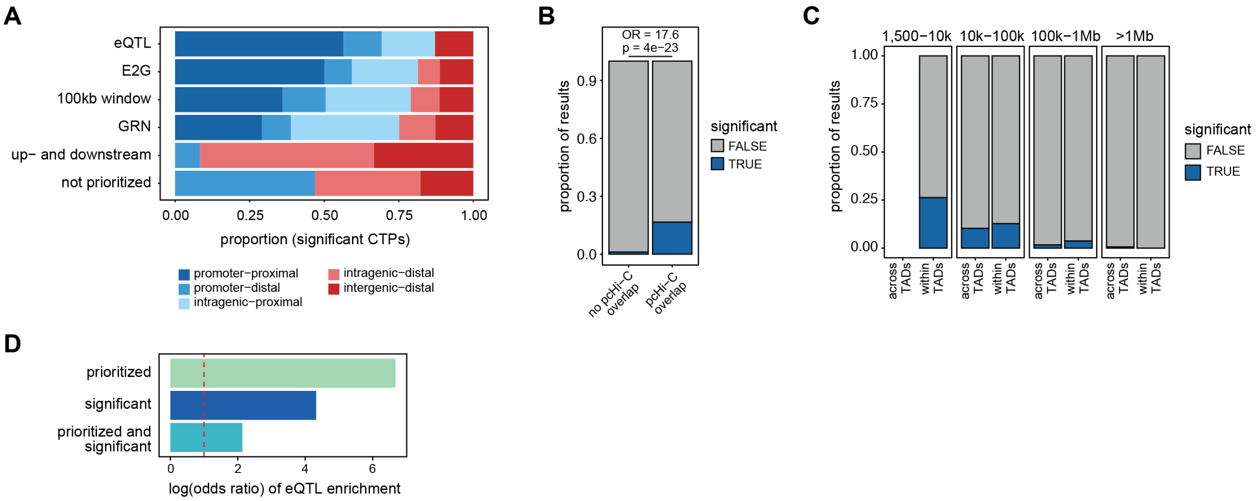


**Fig. S5: Additional results of the CRE screen chromatin analysis.** (**A**) Proportion of significant CRE-gene pairs identified by each of the five CRE-gene prediction strategies and split by the five CRE-gene interaction types. (**B**) Comparison of the proportion of enhancer-like CRE-gene links that are significant if the gene overlaps a promoter and the CRE overlaps a bait in promoter capture Hi-C data (pcHi-C overlap) versus if the gene overlaps a promoter but the CRE does not overlap a bait (no pcHi-C overlap). Odds ratio (OR) and p-value from Fisher exact test of enrichment. (**C**) Comparison of the proportion of enhancer-like CRE-gene links that are significant if the CRE and gene are located within the same topologically associated domain (within TADs), versus if the CRE and gene have a TAD boundary in between them (across TADs). Stratified by CRE-target gene distance. (**D**) Enrichment of whole blood eQTLs among CRE-gene links that were prioritized (top), significant (middle) and both prioritized and significant (bottom), the latter accounting for the fact that the prioritization included eQTL information.

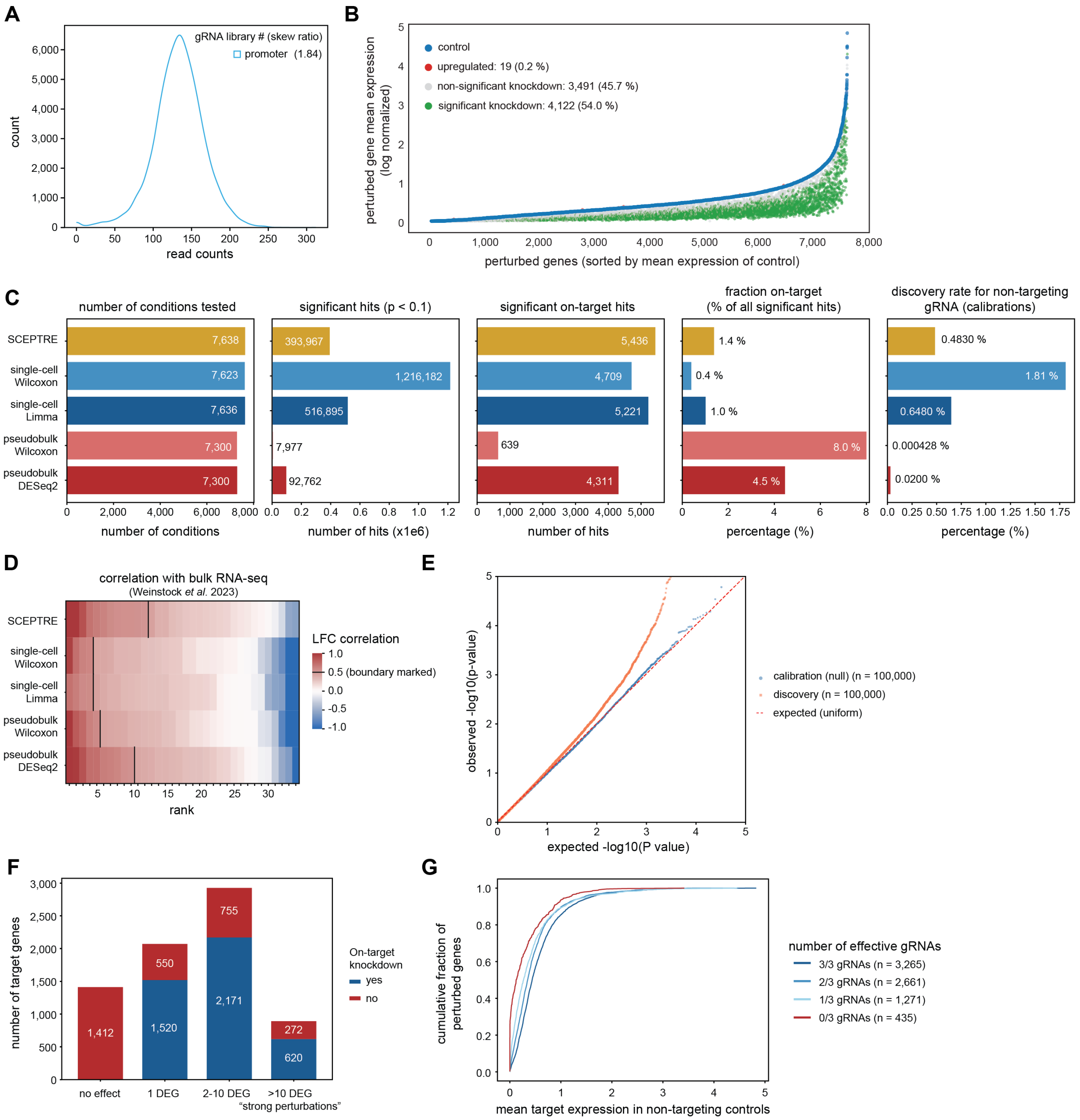


**Fig. S6: Promoter screen analysis additional results.** (**A**) Normalized gRNA count distribution of the promoter screen gRNA library. Skew ratio statistics for the sub-library are listed in the figure. (**B**) On-target knockdown efficiency quantified by contrasting the mean log-normalized expression of the perturbed gene in targeted cells with the mean expression of the same gene in non-targeting control cells. Cells with significant on-target knockdowns identified by DESeq2 are highlighted in green. (**C**) Summary comparison of differential testing tools, including SCEPTRE, limma, Wilcoxon rank-sum tests at the single-cell and pseudobulk levels, and DESeq2. (**D**) Concordance of perturbation effect sizes between single-cell differential expression methods and bulk RNA-seq[40]. For each of five differential testing methods we computed the per-perturbation Pearson correlation between log2 fold changes for all significant genes significant in the bulk data (adjusted $p < 0.05$). Perturbations are ranked by descending correlation independently within each method. (**E**) Quantile–quantile plots of p-value distributions. Observed versus expected −log10(p-values) for DESeq2 calibration (blue) and discovery (red) analyses, subsampled to 100,000 tests. The vertical line indicates the null hypothesis. (**F**) Classification of perturbational effect strength by the number of significant downstream DEGs identified by DESeq2 (adjusted $P < 0.1$), stratified by whether the target gene shows a significant on-target knockdown (blue) or not (red). (**G**) Cumulative distribution of gene expression changes in non-targeting (NT) control cells, grouped by the number of active gRNAs targeting each gene (0 of 2 gRNAs, brown; 1 of 2 gRNAs, yellow; 2 of 2 gRNAs, green).

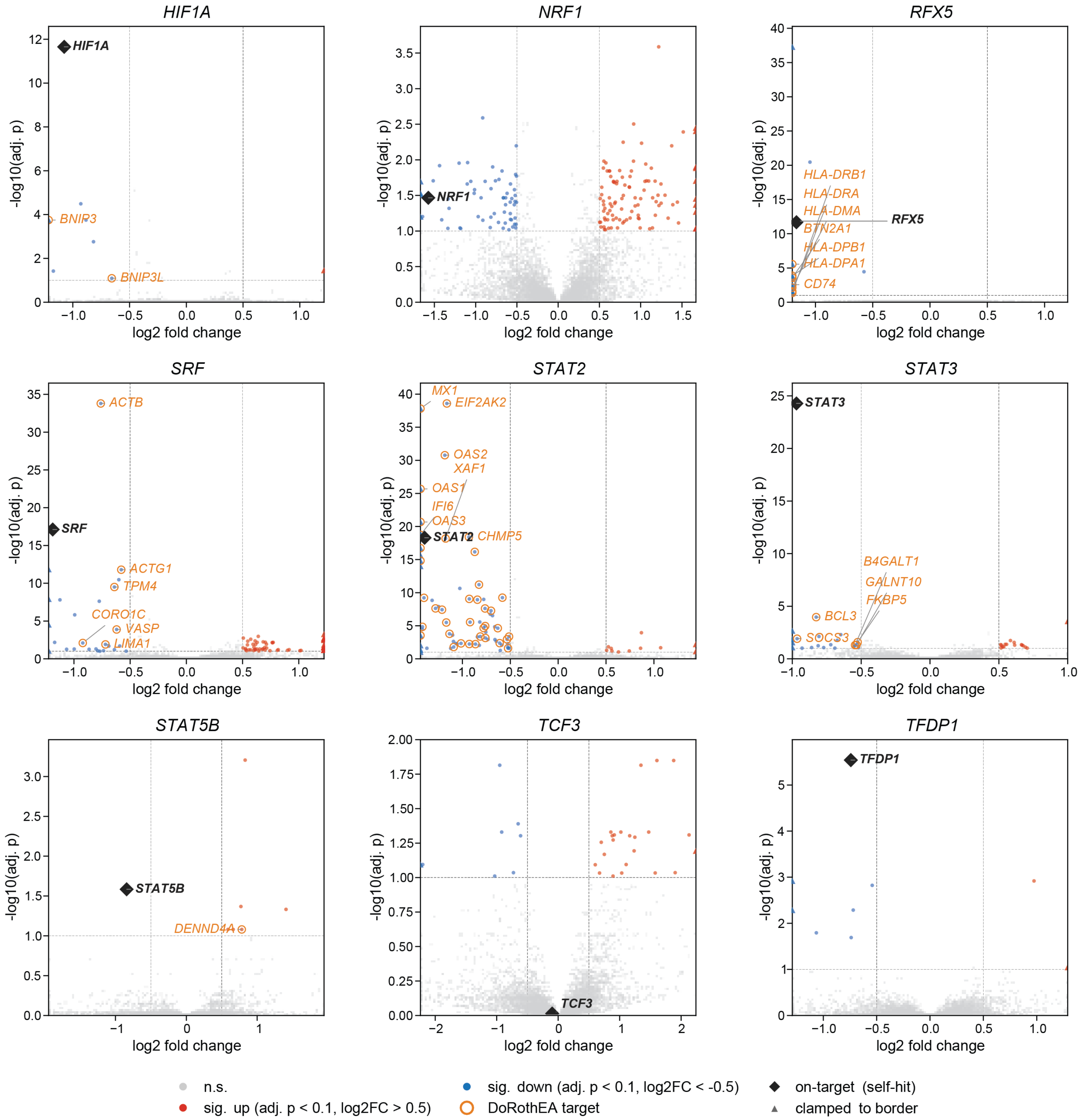


**Fig. S7: Differentially expressed genes and DoRothEA target overlay of perturbed transcription factors.** Volcano plots of the same TFs as shown in Fig. 4E, with differentially expressed genes upon CRISPRi as measured with Perturb-seq. Blue, upregulated; red, downregulated; gray, not significant; orange mark, DoRothEA targets (levels A–C) overlapping with DEGs; diamonds, on-target knockdown. Dashed lines indicate significance cutoffs (|log2FC| > 0.5, adj. p < 0.1).

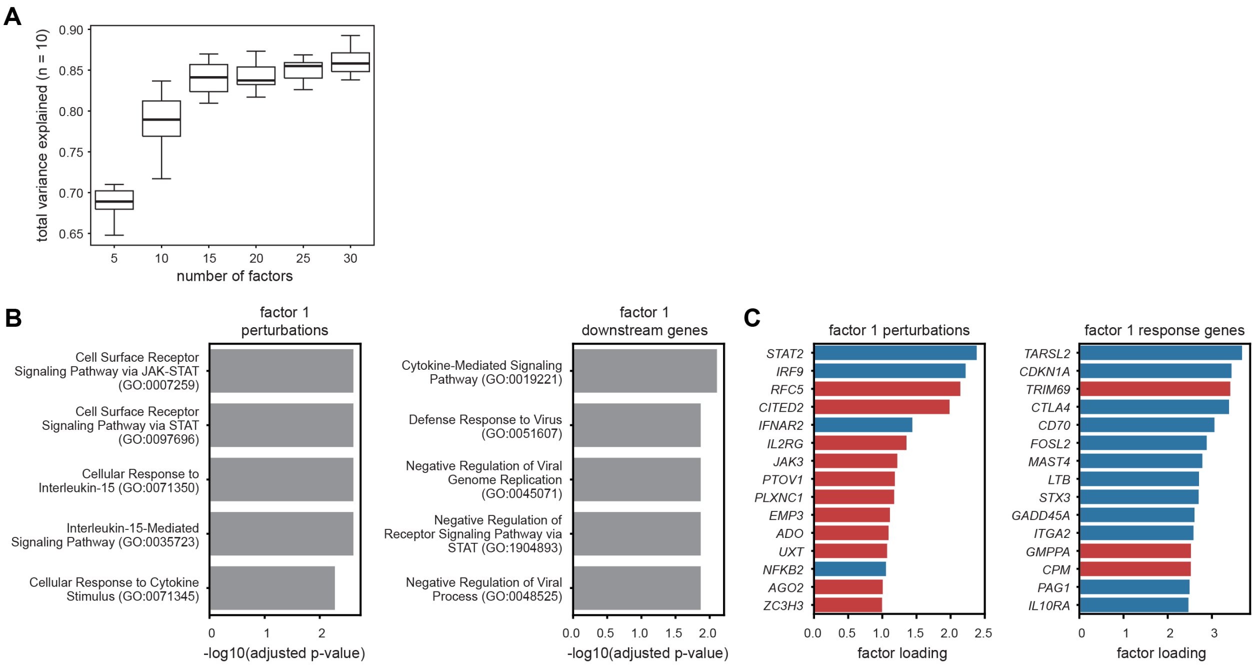


**Fig. S8: Additional results MOFA analysis.** (**A**) Total variance explained when using different numbers of factors in MOFA. For each number of factors, 10 models with different seeds were trained and the total variance explained was calculated as the summed variance explained across all factors. (**B**) Enriched pathways in Factor 1 for the top 10% perturbations (left) and response genes (right). The top 5 most significantly enriched pathways were selected and ordered based on the adjusted p-value. (**C**) Top 15 perturbations (left) and response genes (right) in Factor 1. From the two MOFA matrices (Fig. 4F), the top 15 genes with highest absolute values in Factor 1 were selected and their absolute values displayed. Genes with a negative loading are shown in blue, genes with a positive loading in red.

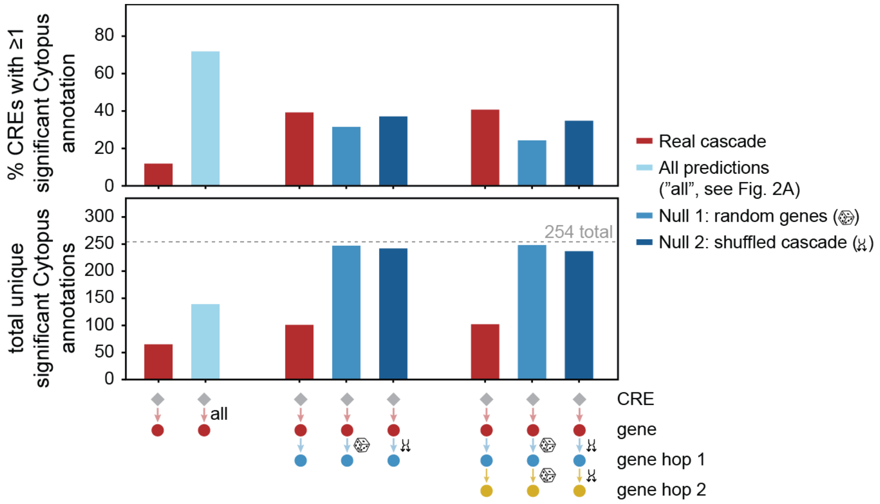


**Fig. S9: Combining CRE screen and promoter screen results.** Percentage of enhancer-like CREs with at least one significant immune cell-type signature from Cytopus v1.3 library (254 terms) [92]. Cytopus was chosen over GO for this analysis since Cytopus terms contain non-overlapping gene sets. Three cascade depths are compared: CRE-to-gene, CRE-to-gene plus one Perturb-seq hop, and CRE-to-gene plus two hops. TAP-seq (red) uses all enhancer-like CRE–gene links; all computational predictions (Fig. 2A) (light blue) serve as TAP-seq null. Perturb-seq nulls: Null 1 (blue), size-matched random genes; Null 2 (dark blue), shuffled cascades (100 permutations each). Upper panel shows the percentage of CREs with Cytopus hits (hypergeometric test, BH-adjusted $p < 0.05$), lower panel shows the total unique Cytopus signatures across all CREs.

**Tables S1-S16**

Table S1: CD4+ T cell donor IDs and metadata

Table S2: GWAS summary statistics

Table S3: Prioritized CRE-gene list

Table S4: ENCODE-rE2G predictions

Table S5: CRE screen TAP-seq primer panel sequences

Table S6: CRE screen gRNA sequences

Table S7: CRE screen TAP-seq primer sequences

Table S8: Pilot library gRNA sequences

Table S9: CRE screen hit list *(will become available upon publication)*

Table S10: eQTL catalogue studies

Table S11: Promoter screen gRNA sequences

Table S12: Promoter screen sequencing summary statistics

Table S13: Promoter screen hit list *(will become available upon publication)*

Table S14: Gene-pairs supported by promoter screen & cis-trans links

Table S15: MOFA clusters

Table S16: Enrichment per disease
